# Functional Plasticity and Evolutionary Adaptation of Allosteric Regulation

**DOI:** 10.1101/2020.02.10.942417

**Authors:** Megan Leander, Yuchen Yuan, Anthony Meger, Qiang Cui, Srivatsan Raman

**Affiliations:** Department of Biochemistry, University of Wisconsin-Madison, Madison, WI – 53706, United States; Department of Bacteriology, University of Wisconsin-Madison, Madison, WI – 53706, United States; Department of Chemistry, Boston University, Boston, MA – 02215, United States

**Keywords:** allostery, deep mutational scanning, functional plasticity

## Abstract

Allostery is a fundamental regulatory mechanism of protein function. Despite notable advances, understanding the molecular determinants of allostery remains an elusive goal. Our current knowledge of allostery is principally shaped by a structure-centric view which makes it difficult to understand the decentralized character of allostery. We present a function-centric approach using deep mutational scanning to elucidate the molecular basis and underlying functional landscape of allostery. We show that allosteric signaling exhibits a high-degree of functional plasticity and redundancy through myriad mutational pathways. Residues critical for allosteric signaling are surprisingly poorly conserved while those required for structural integrity are highly conserved, suggesting evolutionary pressure to preserve fold over function. Our results suggest multiple solutions to the thermodynamic conditions of cooperativity, in contrast to the common view of a finely-tuned allosteric residue network maintained under selection.

## Introduction

Cellular processes are mediated by inter- and intramolecular interactions of proteins. Allostery is intramolecular modulation of protein activity through perturbation at distal sites, and constitutes a dominant mode of post-translational regulation of proteins. Over the decades, we have made major strides in gaining an atomic-level understanding of how proteins fold, catalyze reactions, and interact with other biomolecules. However, understanding the molecular rules governing allostery, a fundamental property of proteins, remains an elusive goal sixty years after its discovery (*1-3*). The knowledge gap exists because decentralized character of allostery makes it challenging to intuitively understand and predict how a distal residue affects an active site some 40-50 Å away (*4*). Mechanisms of catalysis or binding are routinely explained by mutating a limited set of residues as they are driven by local interactions. This classical reductionist approach of studying a limited number of mutations does not scale for a systemic, protein-wide property like allostery as it explores only a small fraction of available sequence space. Therefore, our current understanding of allostery is principally shaped by a structure-centric paradigm based either on conformational heterogeneity (induced fit and conformational selection) (*5-8*), comparison of crystallographic snapshots to infer residues linking allosteric and active sites (*9*), mapping residues undergoing correlated motion by NMR (*10, 11*), or identifying co-evolving residues (*12*). In rare instances, when functional screens were painstakingly carried out, they revealed complex allosteric networks that cannot be gleaned by examining the structure alone (*13*). Therefore, while structure offers vital clues, validating the functional contribution of a residue is the clearest evidence of its role in allostery.

Here, we radically reframe the problem by advancing a function-centric approach to elucidate the molecular basis and functional landscape of allostery. Allosteric switchability is defined as the ability to switch between inactive and active states in a ligand-dependent manner. To investigate the underlying functional landscape, we disrupted allosteric switchability of a bacterial transcription factor (TF) and restored function through alternative paths by systematic, protein-wide mutational scanning. This revealed remarkable functional plasticity as allosteric switchability could be reconstituted time and again after disruption through myriad mutational combinations. While the degree of functional plasticity is site-specific, structural models indicate that recovery of function may be commonly achieved through modulation of DNA or ligand interactions. Phylogenetic analysis revealed that residues critical for allosteric signaling are surprisingly poorly conserved while those required for structural integrity are highly conserved. This suggests stronger evolutionary pressure to preserve fold over function. Molecular dynamics simulations showed conformational states of wildtype are distinct from those of a disrupted mutant, but strikingly similar to a rescued mutant, suggesting different mutational paths lead to the same functional state. Our comprehensive function-centric framework is applicable to any allosteric protein and can lead to biochemical understanding of disease-associated mutations, discovery of druggable allosteric sites, and broad molecular principles of allostery.

### Plasticity of allostery

Our model system is tetracycline repressor (TetR, 207 residues), an all helical (α1-α9), dimeric bacterial TF comprised of ligand- and DNA-binding domains (LBD and DBD) (Fig. S1). As with all allosteric proteins, inactive and active states of TetR correspond to distinct free energy minima (*14*). TetR represses gene expression by binding to promoter (inactive) and ligand induction releases TetR from promoter (active) resulting in transcription. In simplified terms, allosteric switching occurs because free energy of ligand-binding is greater than the free energy difference between inactive and active states, which provides the necessary driving force for conformational stabilization (Fig. 1A, ΔG_LIG_ > ΔG_DIFF,WT_) (*15*). Detailed thermodynamics are described in Supplementary Text (Fig. S2). A mutated TetR may no longer be ligand-inducible if the mutation, without reducing ligand affinity, increases free energy difference (ΔG_DIFF_) between inactive and active states by stabilizing the inactive state, destabilizing the active state, or both. These variants ‘locked’ in a constitutively inactive state are termed ‘dead’ (Fig. 1A, ΔG_LIG_ <ΔG_DIFF,D_). A dead variant may be rescued by a compensatory mutation(s) that restores wildtype-like free energy difference (Fig. 1A, ΔG_LIG_ > ΔG_R_), i.e., ‘rescued’ variants.

**Fig. 1.**
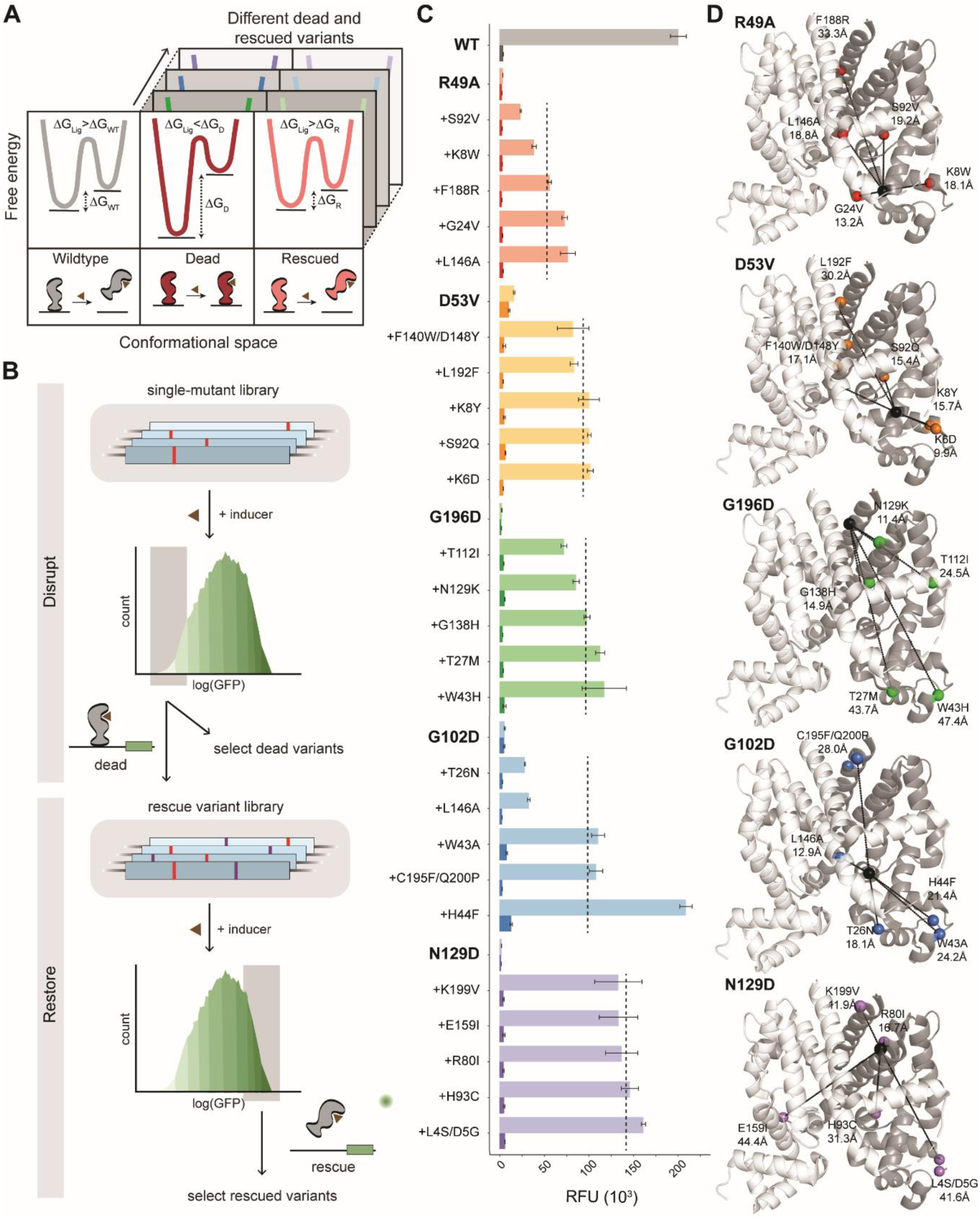
‘Disrupt-and-restore’ strategy to characterize plasticity of TetR allostery. **(A)** A simplified thermodynamic model of allosteric signaling (detailed model Fig. S2). Ligand-binding provides the driving force to switch from inactive to active state in wildtype (left). A disrupting mutation (dead variant) makes TetR constitutively inactive (middle) due to larger energy difference. Allosteric function is restored (rescued variant) by energetic rebalancing via distal compensatory mutations (right). **(B)** Two-stage protein-wide mutational scanning to first disrupt and subsequently restore allosteric signaling using a GFP-based transcriptional reporter system. **(C)** Mean uninduced (light bar) and induced (dark bar) fluorescence of individual variants with the average induced fluorescence of rescued variants indicated (dashed lines). Error bars are standard error of three biological replicates. **(D)** Five dead variants (black spheres), their corresponding rescued variants (colored spheres), and distances between them (dashed lines).

To characterize the plasticity of allosteric networks, we devised a “disrupt-and-restore” strategy. This two-stage, high-throughput, GFP-based mutational screen (*16*) of TetR involves first disrupting and subsequently restoring allosteric signaling (Fig. 1B). We created a library of point mutants by single-site saturation mutagenesis of each TetR residue (207 residues x 19 mutants/residue ≈ 3900 variants) using chip-DNA. First, we sought to disrupt signaling by screening for dead variants. We sorted cells expressing low levels of GFP when incubated with ligand (anhydrotetracycline), and clonally confirmed that the variants are not ligand-inducible. Their ability to repress transcription confirms the dead variants are well-folded proteins. After excluding mutations at ligand-contacting residues, we found allosterically-inactivating dead variants distributed all across TetR, including one (G102D) previously identified 17(*17*). To avoid regional biases in the location of dead variants, we chose five dead variants from different regions for the subsequent rescue screen: R49A (α4 near DBD), D53V (α4 near LBD), G102D (α6 at LBD-DBD interface), N129D (surface exposed on α8), and G196D (α9 at dimerization interface) (Fig. S1). Next, on each dead variant’s background, we constructed another protein-wide single-site saturation mutant library and screened for potential rescued variants by sorting cells expressing high levels of GFP after ligand induction (Fig. 1B). Clonal testing confirmed that allosteric switchability was indeed restored by compensatory mutations in all five dead backgrounds (Fig. 1C, Fig. S3). Compensatory mutations were distributed throughout the protein, some within 10-20 Å and others as far as 40-50 Å away from the site of mutation in the dead variant (Fig. 1D). We compared allosteric activity of rescued variants with wildtype and found that induced reporter fluorescence of rescued variants varied from 10-100% wildtype levels indicating partial to full rescue of function (Fig. 1C). Next, we determined if site-specific bias existed in rescue of function i.e., is the extent of rescue systematically different depending on the dead variant? Site-specific bias was evident in the gradient of mean induced fluorescence of all five rescue variants for each dead variant (Fig 1C, dashed line).

Several important insights emerged from these results. First, TetR exhibits high degree of allosteric plasticity evidenced by the ease of disrupting and restoring function through several mutational paths. This suggests functional landscape of allostery is dense with fitness peaks, unlike binding or catalysis where fitness peaks are sparse. Second, allosterically-coupled residues may not lie along the shortest path linking allosteric and active sites but can occur over long distances because thermodynamic balancing does not require spatial connectivity. Third, allostery signaling occurs through redundant and robust networks instead of a finely-tuned unique pathway.

### Site-specific rescuability of allosteric dysfunction

Two key questions emerged from the screen. Are some dead variants easier to rescue than others and why? Are there common structural mechanisms of rescue? To answer these questions, we sought to exhaustively map and quantify rescuing mutations for each dead variant, and examine possible common rescue modes. We define “rescuability” as the ease of rescuing function and quantify rescuability using the strength of allosteric response and the number of unique rescuing mutations. We sorted each rescue variant library for low GFP cells without inducer to enrich repression-competent variants. For each library, we then clonally induced ∼200 variants and determined the number of unique ligand-responsive clones and their induction level. Relative to wildtype fold induction (45-fold), we classified rescued clones as inactive (<2 fold), moderate (2-10 fold), or strong rescues (10+ fold) (Fig. S3). A clear gradient in rescuability existed from easy to hard as follows: N129D > D53V/G196D > G196D/D53V > R49A > G102D (Fig. 2A). The order of D53V and G196D changed depending on whether fold induction or number of rescuing variants was chosen as the metric of assessment. Leveraging the wealth of dead-rescue allosteric coupling data, we wanted to infer potential structural mechanisms leading to reconstitution of function. We exhaustively mapped each dead variant and their corresponding rescuing mutations to look for focal regions of rescue (Fig. 2B). We observed two broad types of rescuing patterns: specific to individual dead variant and those common to multiple dead variants. The first group were unique rescuing mutations for each dead variant converging at a few key regions, indicating variant-specific regional bias (Fig. 2B, Table S1). However, no obvious structural mechanism of rescue could be gleaned based on the regions where these rescue mutations were found. An exception was N129D for which a high concentration of rescuing mutations fell along α9, suggesting restoration of allosteric signaling through the dimerization interface. The second, more striking group, were rescuing mutations for multiple dead variants converging on specific residues within the the LBD or the DBD. This tantalizingly suggested a common structural mechanism of rescue irrespective of the dead variant. Using Rosetta software, we generated structural models of rescuing mutations at these sites for closer structural inspection. Rescuing mutations at the DBD appeared to generally weaken protein-DNA interactions. Structural model of E37F showed loss of key amino acid-nucleobase interaction and mutation of capping glycine G35 which will likely reduce helix-turn-helix stability (Fig. 2C). In contrast, LBD mutants appeared to strengthen interactions with ligand as seen by additional hydrogen bonds of F67Q and T112Q with the ligand (Fig. 2C). Taken together, these results suggest that rescue of function may be possible through many structural mechanisms. However dominant modes of rescue appear to be mediated through modulation of ligand, DNA, and dimerization interface interactions. Furthermore, we posit that the ease of rescuing a dead variant may correlate with the degree of stabilization of the inactive state (Fig. 2D) i.e., a more stable dead variant could be harder to rescue than a less stable dead variant. This argument is consistent with N129D and G196D being relatively exposed residues (smaller energy perturbation) and hence easier to rescue while the remaining three are more buried and consequently harder to rescue (Fig. 2A).

**Fig. 2.**
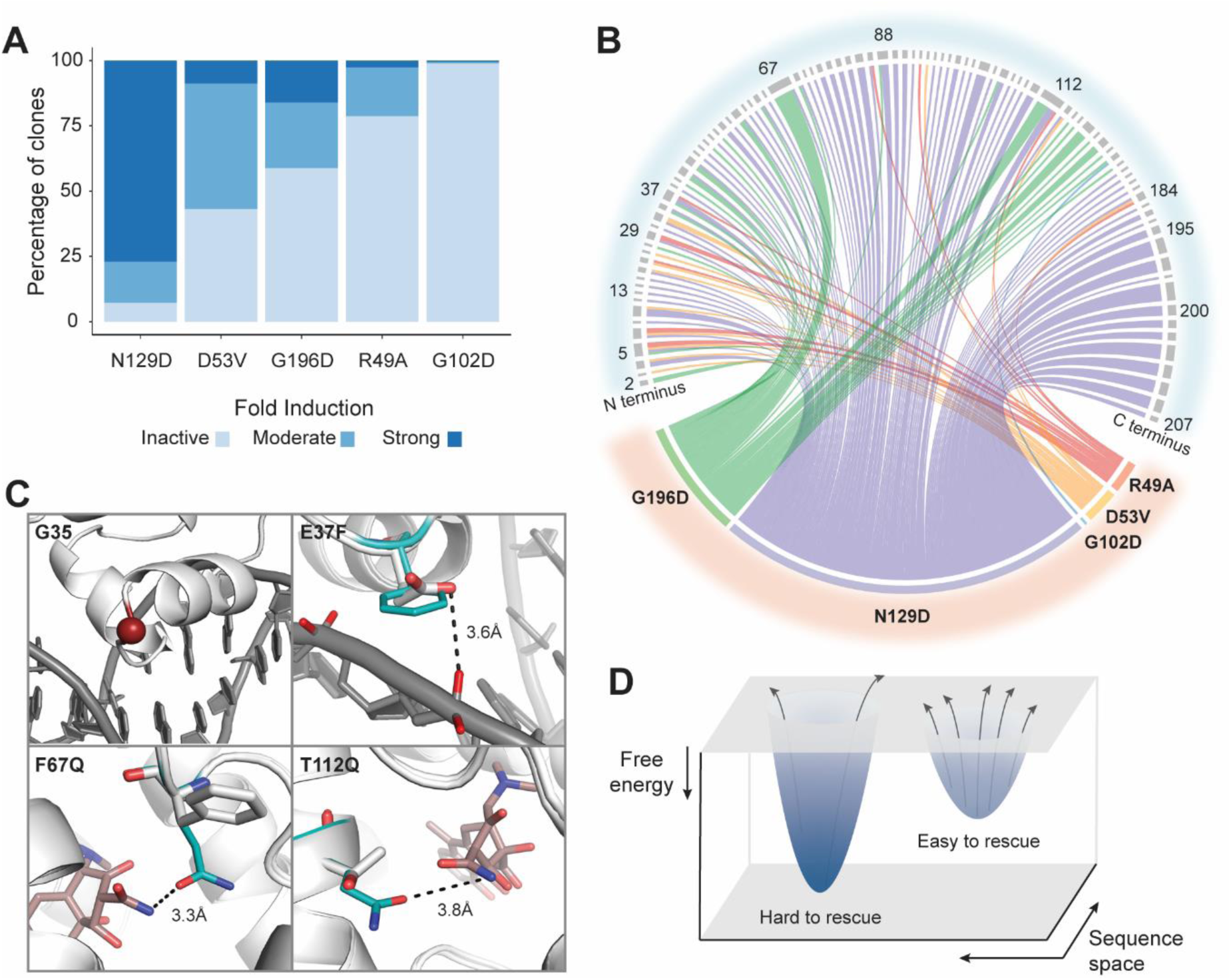
Site-specific differences and structural mechanisms of rescue of function. **(A)** Percentage of rescued clones whose allosteric activity, based on fold induction, is strong (>10), moderate (2-10), or inactive (<2). **(B)** Distribution of rescuing mutations (top) for each dead variant (bottom) shown in different colors. Each line connects dead and rescuing variants and its thickness represents number of rescuing mutations at that position. **(C)** Rosetta structural models of LBD and DBD rescue mutations. Potential common mechanisms of rescue include weaker interaction between DBD and DNA (G35 and E37F) or stronger interactions between LBD and ligand (F67Q and T112Q). **(D)** Free energy landscape hypothesizing relationship between rescuability and energy gap between inactive and active states. Larger gap (deeper well) is harder to rescue than smaller gap (shallower well).

### Structural hotspots more conserved than allosteric hotspots

Next, we examined evolutionary conservation and structural context of residues critical for allosteric signaling (hotspots). To comprehensively map allosteric hotspots, we deep sequenced single-site TetR mutants that repressed GFP, but were not ligand-inducible (Table S2, Data S1). We found hundreds of novel inactivating mutations nearly seven times greater than what was previously known (*13*). After excluding ligand-contacting residues, we classified 55 residues as allosteric hotspots if 5 or more (>25%) mutations at that position inactivated function. The hotspots clustered into four regions. Region 1 is at the interface of the DBD and LBD on α4 and α6, region 2 is a short motif connecting α7 and α8, region 3 is at the dimer interface on α8, and region 4 is at the C-terminal end on α9 (Fig. 3B). Earlier studies identified hotspots in region 1 though not comprehensively, but hotspots in regions 2, 3, and 4 were previously unknown.

**Fig. 3.**
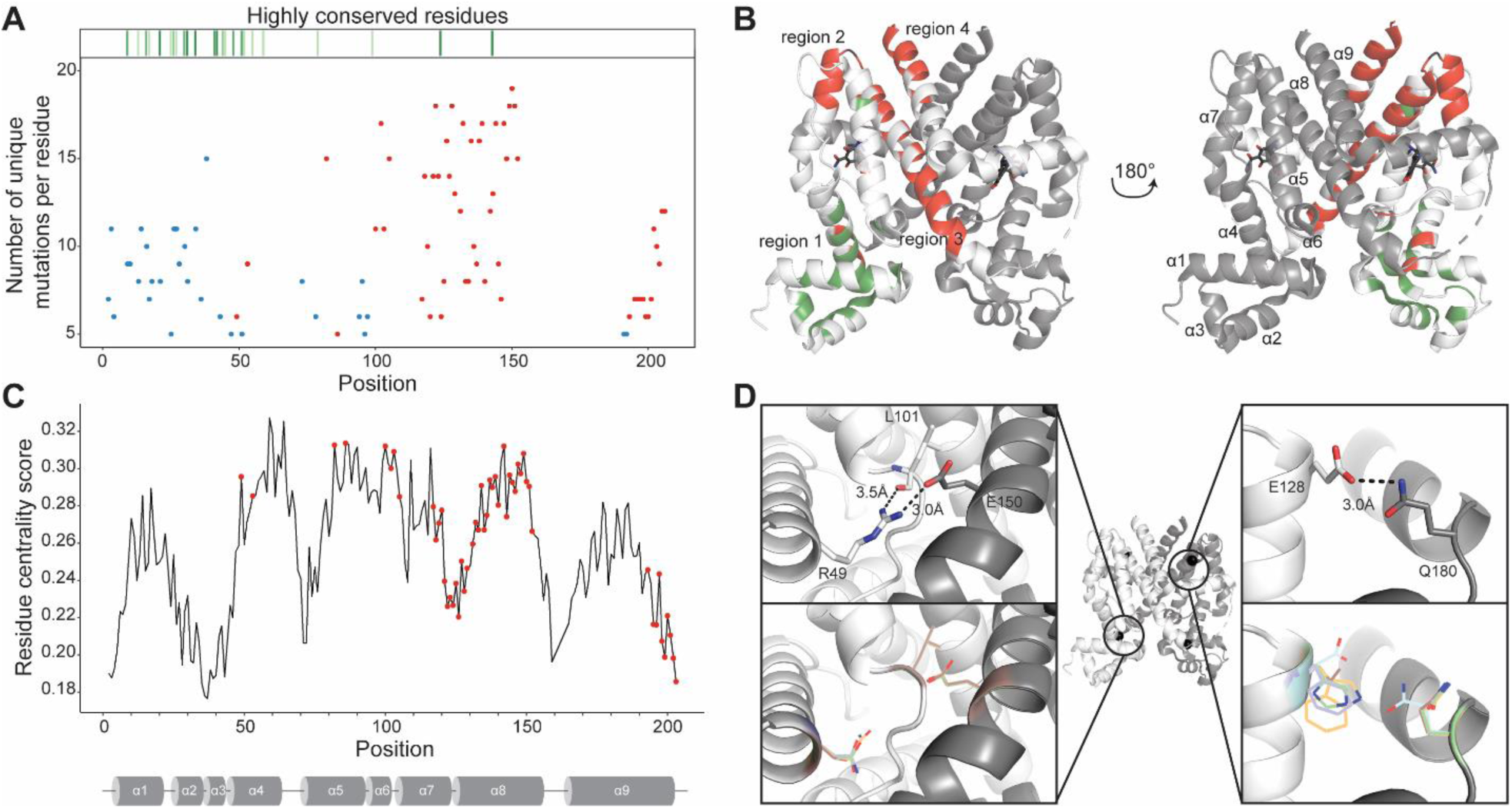
Allosteric hotspots are not highly conserved. **(A)** Comparison of allosteric hotspots (red dot) with highly conserved residues (green lines). Residues critical for structural integrity and activity (DNA-binding) by experimental validation are shown as blue dot. Each dot represents the number of unique point mutants at a position that make TetR constitutively inactive (red) or active (blue). **(B)** Allosteric hotspots (red) and conserved residues (green) separate into distinct groups on TetR structure with only two overlapping residues (black). **(C)** Location of allosteric hotspots (red) overlaid on residue centrality score of every residue (black line). **(D)** Loss of allosteric signaling at hotspots R49 (left) and E128 (right) is likely due to loss of salt bridge or hydrogen bond interactions with inactive mutations.

Catalytic or binding sites are under strong evolutionary selection as reflected in their high sequence conservation even among distant homologs. We aligned TetR-family homologs to determine if allosteric hotspots too are under evolutionary selection. To our surprise, allosteric hotspots showed low sequence conservation in the TetR family. In fact, highly conserved residues and allosteric hotspots neatly separated into distinct, non-overlapping groups (Fig 3A, red points and green lines). Based on their location in the DBD (Fig. 3B, green regions), we hypothesized that conserved residues may be required for structural integrity and DNA binding. Since TetR is a repressor, any mutation that destabilizes or impairs DNA-binding will constitutively express GFP. To evaluate the functional role of conserved residues, we used deep sequencing to determine single-site mutants constitutively expressing reporter without ligand (Fig. S4, Data S1). Indeed, the location of these mutants overlapped with conserved sites confirming the role of conserved residues in maintaining structural integrity and activity (Fig 3A, blue points).

We investigated if allosteric hotspots could be recognized based on their local structural context. Computational approaches predict allosteric hotspots based on the number of contacts a residue makes (residue centrality score) which is rooted in the premise that residues with dense interactions are critical for signal propagation (*18, 19*). We compared residue centrality score at every position and location of allosteric hotspots and found that although some hotspots were correctly identified, the false discovery rate was high (Fig. 3C). Residues that are not allosteric hotspots have high scores (false positives) and conversely those that are have low score (false negatives). Thus, residue centrality captures allosteric signaling through densely-connected interior residues, but misses surface-exposed residues which are often source or sink of allosteric signals. To understand at an atomic level what makes a residue an allosteric hotspot, we modeled using Rosetta two hotspots with high (R49) and low (E128) residue centrality scores (*20*). R49 of one monomer makes a critical salt bridge with D150 in the second monomer. Allosteric signaling may be lost because mutations at this position break the salt bridge (Fig. 3D). Surface exposed residue E128 forms a hydrogen bond with Q180 in α9 of the second monomer. Mutations at E128 lose the hydrogen bond potentially disrupting signaling between dimers (Fig. 3D).

Low conservation of allosteric hotspots, though surprising at first, is consistent with high functional plasticity. Evolution appears to favor preserving structural integrity and activity because their disruption by mutational drift may be harder to restore than allostery. Clustering of hotspots in distinct regions suggests that signaling occurs through coordinated, decentralized action, instead of the shortest path between LBD and DBD. Separation of allosteric hotspots from structural and DBD residues indicates each property could evolve independently leading to orthologs adapted to new niches.

### Free energy landscapes of allosteric mutants

To further investigate if functional similarities between wildtype and rescued variant, and functional differences between wildtype and dead variant are reflected in global structural properties and conformational distributions, we conducted explicit solvent molecular dynamics (MD) simulations at the microsecond time scale. The simulations were carried out for ligand-bound TetR (without DNA) to understand how remote mutations impact the distribution of active and inactive conformations in the ligand-bound state for all three TetR systems (wildtype, dead and rescued). We chose wildtype, G102D (dead) and G102D/C195F/Q200P (rescued) for MD simulations as G102D was hardest to rescue (Fig. 2A). All three TetR systems are structurally stable with a similar backbone RMSD of 2-2.9 Å relative to wildtype crystal structure and similar magnitudes of thermal fluctuations (Fig. 4A). Compared to G102D, the triplet mutant features larger displacements relative to wildtype near the additional mutation sites (C195F/Q200P), including a large fraction of the α4/α5 loop (Fig. 4B). The region in the DNA binding domain that connects α3/α4 also undergoes a larger average displacement relative to wildtype in the triplet mutant compared to G102D (Fig. 4B). Evidently, the additional remote mutations (C195F/Q200P) lead to subtle but persistent structural differences in the DNA binding site more than 35 Å away. We have also examined a range of properties relevant to the motional coupling between protein residues (Fig. S5 and S6), such as positional covariance matrix (Fig. S7), conformational entropy (Fig. S8), distribution of community and hub residues (Fig. S9) and sub-optimal pathways that connect ligand- and DNA-binding sites (Fig. S10). These properties do not exhibit any robust or distinct features among the systems simulated here at the microsecond time scale consistent with our experimental finding that allostery in TetR does not involve a unique, finely-tuned pathway.

**Fig. 4.**
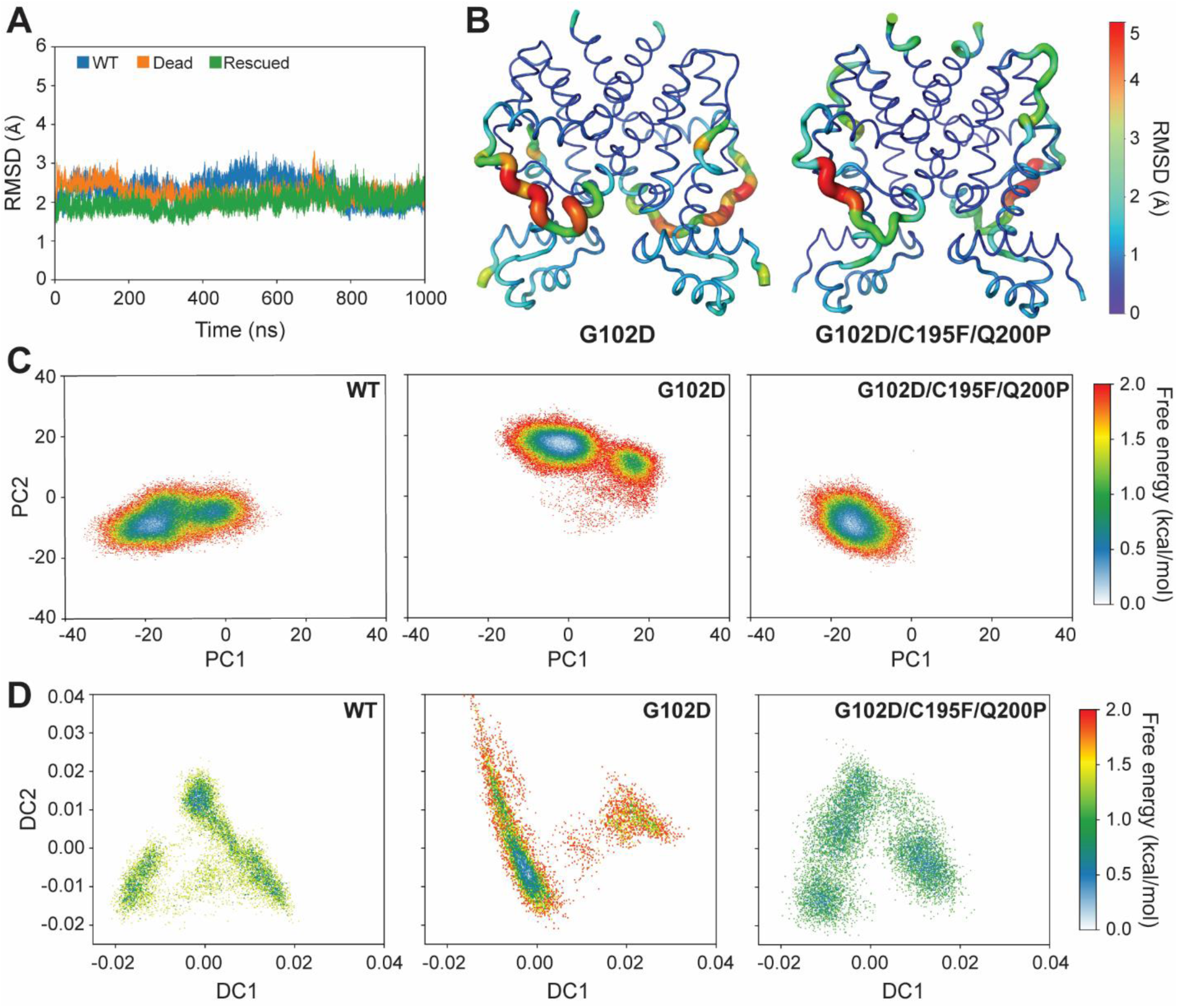
Free energy landscapes of wildtype, dead and rescued mutants consistent with function. **(A)** Backbone RMSDs of wildtype TetR, dead (G102D), and rescued (G102D/C195F/Q200P) variants over time shows stable structures. **(B)** Average structures of dead and rescued variants over the last 800 ns of simulation. Thickness of ribbon and color scale reflect RMSD per residue relative to average wildtype TetR structure over this time window. **(C)** Two-dimensional free energy landscapes projected onto first and second principal components show strong similar conformational distributions of wildtype and rescued variant, but distinct from dead variant. **(D)** Two-dimensional free energy landscapes from the locally scaled diffusion map analysis of WT, dead, and rescued variants also observe similar conformational distributions for the wildtype and rescued variant, with the dead variant being distinct.

However, the underlying free energy landscapes projected onto principal components and locally-scaled diffusion maps (LSDMaps) (*21*) revealed remarkable differences among the three TetR systems (Fig. 4C and 4D). The principal components describe large amplitude motions while the LSDMaps aim to capture slow motions, thus the two types of analyses complement each other (*22*). The free energy landscape projected onto the first two principal components (Fig. 4C, see Fig. S11 for projections along individual principal components) showed a higher degree of similarity between wildtype and the triple mutant, with G102D being clearly different. Similarly, the LSDMap landscapes of the wild type and the rescued mutant resemble each other in shape and locations of free energy basins, while the landscape of the dead mutant is significantly different (Fig. 4D). Projections along higher principal components or diffusion coordinates, by contrast, are similar for all three systems. The first two principal component eigenvectors (Fig. S12 and S13), especially the second one, involve pendulum type of motions of DNA binding domains that were proposed to affect DNA binding affinity (Movie S1 and S2) (*23, 24*). Therefore, the different conformational distributions along the primary principal components in different TetR variants are likely functionally relevant as observed in NMR studies of a TetR homolog, QacR (*25*). However, we caution that it remains challenging to assign quantitative DNA binding affinities to the different conformations sampled in these simulations. Thus, one should not equate Figs. 4C and 4D with the schematic free energy landscapes shown in Fig. 1A or Fig. S2. Nevertheless, these results clearly highlight that mutations remote from the ligand- and DNA-binding sites may lead to sequence-specific changes in the conformational distribution that ultimately get manifested as significant perturbation in function. It is encouraging that these changes are observed in unbiased MD simulations at the microsecond time scale, and we anticipate that additional insights can be gleaned with enhanced sampling simulations in the future.

## Conclusions

The conclusions of this study have profound impact on evolutionary, structural and biophysical understanding of allostery. High functional plasticity of TetR may facilitate a TetR-like protein to undertake adaptive walks in mutational space to acquire new ligand specificities. This may partly explain the extraordinary diversity of ligands binding TetR-family proteins (*26*). A critical assumption in statistical coupling analysis and other co-evolution-based approaches is that nature imposes evolutionary pressure at specific sites to preserve allosteric coupling. Our case study of TetR challenges this assumption because we find that allosteric sites are not necessarily conserved, though allostery by itself may be conserved (*27*). Conservation of specific sites may be a feature of catalysis or binding, which strongly depend on key local molecular interactions. In contrast, since allostery is systemic, balancing of thermodynamic forces can be satisfied by decentralized functional constraints in many possible ways without the need for conserving specific sites. Future mutational studies on other TetR-family members may reveal if common lynchpin sites and couplings exist within the family, that can be recognized from sequences alone. Together with machine learning, large datasets like ours linking structural effects of mutations to function can help develop heuristic molecular rules, such as gain/loss of salt bridges or hydrogen bonds (R49 and E128, Fig 3D), for allosteric communication (*28*). Furthermore, simulating molecular trajectories of tens or hundreds of mutants will help us interpret how conformational heterogeneity of protein ensembles is related to protein function. This is an exciting new direction for the field of MD simulations where MD is used not just as a validation and explanatory tool, but also for function prediction.

## Materials and Methods

### Plasmid construction

We constructed a sensor plasmid with TetR(B) (Uniprot #P04483) cloned into a low-copy backbone (SC101 origin of replication) carrying spectinomycin resistance. The *tetRb* gene was driven by a variant of promoter apFAB61 and Bba_J61132 RBS (*29*). On a second plasmid, superfolder GFP (*30*) was cloned into a high-copy backbone (ColE1 origin of replication) carrying kanamycin resistance under the control of the ptetO promoter. To control for plasmid copy number, RFP was constitutively expressed with the BBa_J23106 promoter and Plotkin RBS (*29*) in a divergent orientation to sfGFP.

### Library synthesis

A comprehensive single-mutant TetR library was generated by replacing wild-type residues at positions 2-207 of TetR to all other 19 canonical amino acids (3,914 total mutant sequences). Oligonucleotides encoding each single point mutation were synthesized as single-stranded Oligo Pools from Twist Bioscience and organized into subpools spanning the *tetRb* gene: residues 2–39, 40–77, 78–115, 116–153, 154–191, 192–207. Oligo pools were encoded as a concatemer of the forward priming sequence, a BasI restriction site (5’-GGTCTC), six-base upstream constant region, *tetR* mutant sequence, six-base downstream constant region, a BsaI site (5’-GAGACC), and the reverse priming sequence. Subpools were resuspended to 25 ng/μl and amplified using primers specific to each oligonucleotide subpool with KAPA SYBR FAST qPCR (KAPA Biosystems; 1 ng template). A second PCR amplification was performed with KAPA HiFi (KAPA Biosystems; 1 μl qPCR template, 15 cycles maximum). We amplified corresponding regions of pSC101_tetR_specR with primers that linearized the backbone, added a BsaI restriction site, and removed the replaced wild type sequence. Vector backbones were further digested with DpnI, BsaI, and Antarctic phosphatase before library assembly.

We assembled mutant libraries by combining the linearized sensor backbone with each oligo subpool at a molar ratio of 1:5 using Golden Gate Assembly Kit (New England Biolabs; 37 °C for 5 min and 60 °C for 5 min, repeated 30x). Reactions were dialyzed with water on silica membranes (0.025 μm pores) for 1 hour before transformed into DH10β cells (New England Biolab). Library sizes of at least 100,000 CFU were considered successful. DH5α cells (New England Biolab) containing the reporter pColE1_sfGFP_RFP_kanR were transformed with extracted plasmids to obtain libraries of at least 100,000 CFU.

Rescue variant libraries were synthesized as described above using sensor plasmid of each dead variant as the linearized backbone. To avoid mutations close in sequence space, the oligo subpool containing the mutation was not cloned into the library – for example, the G102D library did not contain mutations spanning residues 78-115 in TetR. All double-mutant libraries contained 3,192 possible sequences, except for the G196D rescue library which contained 3,610 sequences.

### Fluorescence activated cell sorting

All library cultures and clonal variants were grown for 16h at 37°C in LB containing 50 ug/mL kanamycin and 50 ug/mL spectinomycin. Libraries were seeded from a 50 µL aliquot of glycerol stocks and grown to an OD_600_ ∼.2 before induction with 1 µM anhydrotetracycline (aTC). Saturated library cultures were diluted 1:50 in 1x PBS and fluorescence intensity was measured on a SH800S Cell Sorter (Sony). Remaining uninduced cultures were spun down and plasmids were extracted for next-generation sequencing (NGS) to represent the presorted library distribution. We first gated cells to remove debris and doublets, and selected for variants constitutively expressing RFP. The induction profile of wild type TetR was used as reference in drawing gates on GFP fluorescence to identify dead and rescued variants. Gates were drawn between either ∼10-1000 RFU (fluorescence distribution of repressed wildtype TetR, dead) or ∼30,000-200,000 RFU (fluorescence distribution of inducted wildtype TetR, rescued) dependent on population. A total of 500,000 events were sorted for each gated population and cells were recovered in 5mL for 1 hour before cultures were plated to clonally screen for inactive or rescue variants. Antibiotics were added and cultures were grown for an additional 6h until an OD_600_ ∼.2 was reached before plasmid extraction for sequencing.

To identify initial dead variants for further analysis, the TetR single-mutant library was induced with aTC and nonfluorescent cells were sorted. We screened ∼300 colonies from the sorted library and clonally identified dead variants. Similarly, fluorescent cells of the rescue libraries were sorted and ∼200-400 colonies screened to identify rescued variants.

### Clonal screening and flow cytometry

To screen for dead and rescuing variants, individual colonies were picked and grown to saturation in 96-well plates for 8h. Non-fluorescent colonies under blue light were selected in these screens to select variants with DNA-binding capability in the absence of inducer. Saturated cultures were diluted 1:50 in LB-kan/spec and grown in the presence and absence of 1 µM aTC for 16h before OD_600_ and GFP fluorescence (Gain: 40; Excitation: 488/20; Emission: 525/20) were read on a multi-plate reader (Synergy HTX, BioTek). Fluorescence was normalized to OD_600_ and the fold inductions for each variant were computed as the ratio of induced and non-induced fluorescences. Variants with at least 10-fold induction were plated, confirmed in triplicate, and Sanger sequenced (Functional Bioscience). Selected variants were diluted 1:50 into 1x PBS before being measured on the Attune NxT Flow Cytometer (Thermo Fisher Scientific).

### Quantifying rescuability

Rescue variant libraries were grown in the absence of inducer and nonfluorescent cells were sorted to select for DNA-bound variants. For each rescue variant library, we randomly selected and screened 192 nonfluorescent colonies after sorting to determine the percentage of cells that recovered some degree of function. Fold induction for each individual variant was measured on the plate reader and variants with at least 5-fold induction were pooled and plasmids extracted for sequencing. To identify the number of unique variants in the screen, an initial group of all 192 variants was pooled and prepped for sequencing as well.

### Next-generation sequencing (NGS) and analysis

Libraries were prepared from plasmids extracted from the presorted, sorted, and pooled libraries to identify dead and rescued variants. Libraries were sequenced with a 2×250 sequencing run in which the TetR gene was broken into two fragments to cover the entire encoding region. Libraries were amplified with two primer sets, one specific to the encoding region of interest (residues 2-115 or 116-207) adding the NGS priming region, and a second outer pair adding the unique barcode and library adapter. Libraries were amplified in a 2-step PCR reaction. First, inner primers were added at a final concentration of 125nM each and reaction run for 11 cycles before 7x the concentration of the outer stem primers were added (.9uM) and another 8 cycles were run. Inactive variant libraries were sequenced with a 2×250 MiSeq run at the UW-Madison Biotechnology Center and rescuability screens were sequenced with a 2×250 run through Genewiz (Amplicon-EZ).

Paired-end Illumina sequencing reads were merged with FLASH (Fast Length Adjustment of SHort reads) using the default software parameters (*31*). Phred quality scores (Q) were used to compute the total number of expected errors (E) for each merged read (*32*). Reads exceeding the maximum expected error threshold (Emax) of 1 were removed. To identify inactive variants, nonfluorescent cells in the single-mutant library were sorted in duplicate in both the presence and absence of 1 µM aTC and prepped for sequencing along with the initial, unsorted library. Sequencing reads within each barcoded sample were normalized to 200,000 total reads and then by the number of variants within group before a cutoff of 10 reads was applied to reduce noise.

Variants with at least 25 reads in both the presence and absence of ligand in both replicates were identified as ‘dead’. Positions with five or more dead variants were termed allosteric, or dead, hotspots. Variants present in the presorted libraries but not in either nonfluorescent (± 1 µM aTC) were assumed to be largely fluorescent. We termed these variants ‘broken’ as they are predicted to destabilize the protein and/or affect DNA-binding. Positions in which five or more mutations broke the protein were termed broken hotspots. Sequencing of variants from the rescuability screen were prepared as above and a cutoff of 10 reads was applied to identify the presence of a variant. Rescued variants with two or more compensatory mutation were found in the screen, but only single-mutant compensatory mutations were used for further analysis.

### Sequence conservation

TetR homologs were generated using HMM search (https://www.ebi.ac.uk/Tools/hmmer/) (*33*) against UniProtKB database with TetR(B) sequence as query. Sequences with alignment coverage less than 95% of full length TetR(B) were removed from consideration. The remaining sequences were aligned using Clustal Omega (*34*). After applying a redundancy cutoff of 90%, we were left with approximately 1900 sequences which was used to evaluate sequence conservation score within Jalview (*35*). Conservation score in Jalview is computed by AMAS tool (*36*) and positions with a score of seven or more were termed highly conserved.

### ddG calculations and structural models of mutations

The crystal structure of TetR(B) with bound [Minocycline:Mg]^+^ and DNA-bound TetR(D) dimer structures were obtained (PDB ID: 4ac0, 1qpi) and water molecules removed before calculations run; the bound ligand was also removed from TetR(B). All modeling calculations were performed using the Rosetta molecular modeling suite v3.9. Single-point mutants were generated using the standard ddg_monomer application (*20*), which enables local conformational to minimize energy. For TetR(B), calculations were run at every position in protein for all 20 amino acids, generating 50 possible mutant and WT structural models for each protein variant. Structures with the lowest total energy from the 50 mutant and WT models were used to calculate ddG and served as models for structural analysis. Calculations for DNA-bound TetR(D) variants were prepared and analyzed in similarly, though only select rescuing mutations at G35 and E37 were run.

### Molecular dynamics simulations

The missing residues 160-164 of the TetR(B) crystal structure (PDB ID: 4ac0) were modeled in CHARMM-GUI. To ensure a stable hydrogen bond interaction between residue 64 and aTC, the protonation state of residue 64 was set to be HSE, as was done in a previous computational analysis of the TetR system (*37*). For mutants, corresponding residues were mutated.

For each TetR system, the initial structure was solvated in a rectangular box solvated with ∼27,300 TIP3P water molecules with a 15.0 Å of edge distance. 87-89 Na+ and 77 Cl-counter ions were added to ensure neutrality at an ionic strength of 0.15 M, resulting in a box of around 88 × 88 × 88 Å^3^ using periodic boundary conditions. All simulations (∼88,700 total atoms) were performed in OpenMM using the CHARMM36m force field. CHARMM-GUI was used to generate input files. Particle mesh Ewald (PME) with the Ewald error tolerance of 0.0005 was used to calculate electrostatic interactions; the tolerance stands for the average fractional error in forces that is acceptable. The van der Waals interaction was treated with a switching scheme for distances between 10 and 12 Å. Energy minimization was carried out with 10000 steps of the L-BFGS algorithm. The system was then equilibrated in the NVT ensemble at 303.15 K for 250 ps. Langevin dynamics was used with a collision frequency of 1 ps^-1^. During minimization and equilibration, small harmonic restraints were applied to both protein backbone and sidechain atoms, with force constants of 400 kJ/(mol nm^-2^) and 40 kJ/(mol nm^-2^), respectively. After equilibration, all atoms were released and no restraint was applied. Production simulations were carried out in the NPT ensemble at 303.15K using Langevin dynamics with a friction coefficient of 1 ps^-1^. MonteCarloBarostat was used with the pressure of 1 bar and the pressure coupling frequency of 100 steps. In equilibration and production runs, all water molecules were rigid and all bonds involving hydrogen atoms were constrained using HBonds constraints in OpenMM, allowing a time step of 2 fs.

## Acknowledgments

We thank Dr. Elizabeth Craig and Dr. Katherine Henzler-Wildman for critical review of the manuscript.

## Author Contributions

M.L., Y.Y., Q.C., and S.R. conceived the study. M.L. and S.R. designed experiments. M.L. performed experiments. M.L. and S.R. analyzed experimental data. A.M. carried out Rosetta calculations. Y.Y. and Q.C. designed and carried out MD simulations. M.L., Y.Y., Q.C., and S.R. wrote the manuscript.

## Data and Materials Availability

All data is available in the manuscript or the supplementary materials

## References

1. S. J. Wodak et al., Allostery in its many disguises: from theory to applications. Structure 27, 566–578 (2019).

2. J. P. Changeux, S. J. Edelstein, Allosteric mechanisms of signal transduction. Science 308, 1424–1428 (2005).

3. J. P. Changeux, 50 years of allosteric interactions: the twists and turns of the models. Nat. Rev. Mol. Cell Biol. 14, 819–829 (2013).

4. N. V. Dokholyan, Controlling allosteric networks in proteins. Chem. Rev. 116, 6463–6487 (2016).

5. Q. Cui, M. Karplus, Allostery and cooperativity revisited. Protein Sci. 17, 1295–307 (2008).

6. J. Guo, H. X. Zhou, Protein allostery and conformational dynamics. Chem. Rev. 116, 6503–6515 (2016).

7. V. J. Hilser, Biochemistry. An ensemble view of allostery. Science 327, 653–654 (2010).

8. K. Gunasekaran, B. Ma, R. Nussinov, Is allostery an intrinsic property of *all* dynamic proteins? Proteins 57, 433–443 (2004).

9. M. D. Daily, J. J. Gray, Local motions in a benchmark of allosteric proteins. Proteins 67, 385–399 (2007).

10. M. J. Holliday, C. Camilloni, G. S. Armstrong, M. Vendruscolo, E. Z. Elsenmesser, Networks of dynamic allostery regulate enzyme Function. Structure 25, 276–286 (2017).

11. N. Popovych, S. Sun, R. H. Ebright, C. G. Kalodimos, Dynamically driven protein allostery. Nat. Struct. Mol. Biol. 13, 831–838 (2006).

12. G. M. Süel, S. W. Lockless, M. A. Wall, R. Ranganathan, Evolutionarily conserved networks of residues mediate allosteric communication in proteins. Nat. Struct. Biol. 10, 59–69 (2003).

13. G. Müller, B. Hecht, V. Helbl, W. Hinrichs, W. Saenger, W. Hillen, Characterization of non-inducible Tet repressor mutants suggests conformational changes necessary for induction. Nat Struct. Biol. 2, 693–703 (1995).

14. G. Chure, M. Razo-Mejia, N. M. Belliveau, T. Einav, Z. A. Kaczmarek, S. L. Barnes, M. Lewis, R. Phillips, Predictive shifts in free energy couple mutations to their phenotypic consequences. Proc. Natl. Acad. Sci. USA 116, 18275–18284 (2019).

15. C. J. Tsai, R. Nussinov, A unified view of “how allostery works”. PLoS Comput. Biol. 10, e1003394 (2014).

16. D. M. Fowler, S. Fields, Deep mutational scanning: a new style of protein science. Nat. Methods 11, 801–807 (2014).

17. O. Scholz, E. M. Henssler, J. Bail, P. Schubert, J. Bogdanska-Urbaniak, S. Sopp, M. Reich, S. Wisshak, M. Köstner, R. Bertram, W. Hillen, Activity reversal of Tet repressor caused by single amino acid exchanges. Mol. Microbiol. 53, 777–789 (2004).

18. G. Amitai, A. Shemesh, E. Sitbon, M. Shklar, D. Netanely, I. Venger, S. Pietrokovski, Network analysis of protein structures identified functional residues. J. Mol. Biol. 344, 1135–1146 (2004).

19. A. del Sol, H. Fujihashi, D. Amoros, R. Nussinov, Residues crucial for maintaining short paths in network communication mediate signaling in proteins. Mol. Syst. Biol. 2, 2006.0019 (2006).

20. E. Kellogg, A. Leaver-Fay, D. Baker, Role of conformational sampling in computing mutation-induced changes in protein structure and stability. Proteins 79, 830–838 (2011).

21. M. A. Rohrdanz, W. Zheng, M. Maggioni, C. Clementi, Determination of reaction coordinates via locally scaled diffusion map, J. Chem. Phys., 134, 124116 (2011).

22. Y. Zheng, Q. Cui, The histone H3 N-terminal tail: a computational analysis of the free energy landscape and kinetics, Phys. Chem. Chem. Phys. 17, 13689–13698 (2015).

23. A. Aleksandrov, L. Schuldt, W. Hinrichs, T. Simonson, Tet repressor induction by tetracycline: a molecular dynamics, continuum electrostatics, and crystallographic study. J. Mol. Biol. 378, 898–912 (2008).

24. P. Orth, D. Schnappinger, W. Hillen, W. Saenger, W. Hinrichs, Structural basis of gene regulation by the tetracycline inducible Tet repressor-operator system. Nat. Struct. Biol. 7, 215–219 (2000).

25. K. Takeuchi, M. Imai, I. Shimada, Conformational equilibrium defines the variable induction of the multidrug-binding transcriptional repressor QacR. Proc. Natl. Acad. Sci. USA 116, 19963–19972 (2019).

26. L. Cuthbertson, J. R. Nodwell, The TetR family of regulators. Microbiol. Mol. Biol. Rev. 77, 440–475 (2013).

27. A. A. Fodor, R. W. Aldrich, On evolutionary conservation of thermodynamic coupling in proteins. J. Biol. Chem. 279, 19046–19050 (2004).

28. O. N. Demerdash, M. D. Daily, J. C. Mitchell, Structure-based predictive models for allosteric hot spots. PLoS Comput. Biol. 5, e1000531 (2009).

29. S. Kosuri, D. B. Goodman, G. Cambray, V. K. Mutalik, Y. Gao, A. P. Arkin, D. Endy, G. M. Church, Composability of regulatory sequences controlling transcription and translation in *Escherichia coli*. Proc. Natl. Acad. Sci. USA 110, 14024–14029 (2013).

30. J. D. Pédelacq, S. Cabantous, T. Tran, T. C. Terwilliger, G. S. Waldo, Engineering and characterization of a superfolder green fluorescent protein. Nat. Biotechnol. 24, 79–88 (2006).

31. T. Magoč, S. L. Salzberg, FLASH: fast length adjustment of short reads to improve genome assemblies. Bioinformatics 27, 2957–2963 (2011).

32. R. C. Edgar, H. Flyvbjerg, Error filtering, pair assembly and error correction for next-generation sequencing reads. Bioinformatics 31, 3476–3482 (2015).

33. S. C. Potter, A. Luciani, S. R. Eddy, Y. Park, R. Lopez, R. D. Finn, HMMER web server: 2018 update. Nucleic Acids Res. 46, W200–W204 (2018).

34. F. Sievers, A. Wilm, T. J. Gibson, K. Larplus, W. Li, R. Lopez, H. McWilliam, M. Remmert, J. Söding, J. D. Thompson, D. G. Higgins, Fast, scalable generation of high-quality protein multiple sequence alignments using Clustal Omega. Mol. Syst. Biol. 7, (2011).

35. A. M. Waterhouse, J. B. Procter, D. M. Martin, M. Clamp, G. J. Barton, Jalview version 2-a multiple sequence alignment editor and analysis workbench. Bioinformatics 25, 1189–1191 (2009).

36. C. D. Livingstone, G. J. Barton, Protein sequence alignments: a strategy for the hierarchical analysis of residue conservation. Comput. Appl. Biosci. 9, 745–758 (1993).

37. A. Aleksandrov, J. Proft, W. Hinrichs, T. Simonson, Protonation patterns in Tetracycline:Tet Repressor recognition: simulations and experiments. Chem. Bio. Chem. 8, 675–685 (2007).

38. J. Eargle, Z. Luthey-Schulten, NetworkView: 3D display and analysis of protein·RNA interaction networks. Bioinformatics 28, 3000–3001 (2012).

