## Supplemental for "Functional Plasticity and Evolutionary Adaptation of Allosteric Regulation"

**This PDF file includes:**

Supplementary Text
Figures S1 to S13
Tables S1 to S3
Scheme S1
Legends for Movies S1 to S2
Legends for Dataset S1
SI References

**Other supplementary materials for this manuscript include the following:**

Movies S1 to S2
Dataset S1 – All_Mutations_Summary.xlsx

### Supplementary Information Text

**Convergence of different properties in MD simulations.** As shown in Figs. S5 and S6, the average structure converges rather quickly, reflecting the general structural rigidity of the system. For example, the average structures after 100 ns and 1 μs do not exhibit any major difference at the backbone level (Fig. S6) even for the loop regions, which exhibit higher thermal fluctuations compared to the helices (Fig. S5). The DNA binding domains, despite considerable thermal fluctuations, overlap well for the averaged structures over 100 ns and 1 μs timescales. The general patterns for RMS fluctuations also do not show any major variation beyond 100 ns of sampling, except for several loop regions. These trends apply to all the three TetR variants studied here.

By contrast, properties that reflect correlated motions converge much more slowly. For example, the covariance matrices among the three TetR variants appear rather different after 100 ns (Fig. S7D-F). Following 1 μs of sampling, however, the differences become substantially smaller (Fig. S7A-C). The convergence of configurational entropy, which depends on the covariance matrix, is even more difficult, as illustrated by Fig. S8. The overall trend in the computed configurational entropy, which does not clearly distinguish the dead (G102D) and rescuing (G102D/C195F/Q200P) mutants, is similar when only Cα or all heavy (non-hydrogen) atoms are included in the covariance and entropy calculations. These observations highlight the importance of sampling for probing correlated motions in even relatively rigid systems such as TetR.

**Community analysis.** In the analysis of protein allostery, it is common to decompose the protein structure into communities, which feature strong intra-community correlations (*1*, *2*). In Fig. S9, we show the results of community analysis for the three TetR variants following 100 ns and 1 ms of sampling. Evidently, the results of community analysis with different amounts of sampling can change considerably, again highlighting the slow convergence of properties that depend on the covariance matrix. Following 1 μs of sampling, the protein structure is divided into several communities and the general pattern does not differ considerably among the three TetR variants studied here, as expected based on the overall similar covariance matrices (Fig. S7). The DNA binding domain is classified as a single community, and it does not contain a significant number of residues in the ligand binding domain. Therefore, the degree of direct coupling between the DNA binding and ligand binding domains is relatively weak, which suggests that allosteric coupling between them occurs through indirect correlations.

Accordingly, we also performed suboptimal path analysis and identified "hub residues" that mediate the "information flow" between these two functional domains. The "hub residues" are sorted in a descending order by the occurrence of each residue in suboptimal paths. The top 50 most-occurred "hub residues" were selected to compare with the "hotspot" residues found experimentally. As shown in Fig. S10, the common set between the two sets of residues contains only a handful of residues, which constitute about 10% percentage of either set. Evidently, the functional significance of "hub residues", at least in TetR based on microsecond simulations, is limited.

### Materials and Methods

**Centrality scoring.** Centrality scores of each residue were calculated using the Network Analysis of Protein Structures (NAPS) server (http://bioinf.iiit.ac.in/NAPS/) (*4*). The unweighted atom pair contact network of the wildtype TetR(B) dimer (PDB ID: 4ac0) was generated using a 0-5 Å threshold. Node centrality was then measured by closeness, or the shortest distance of one position to all others in the network.

**Modeling of anhydrotetracycline ligand.** The ligand anhydrotetracycline (aTC) was built in Avogadro. To ensure the strong interaction with $Mg^{2+}$, the hydrogen atom on the O2 atom was removed; this was also consistent with findings from previous electrostatics calculations by Simonson and co-workers. A structure with a total charge of -1 was obtained. The molecule was then optimized in Gaussian using B3LYP/6-31G(d). The optimized structure was uploaded to CHARMM-GUI. Atom types and partial charges were assigned based on the CHARMM General Force Field (CGenFF). The obtained structure and force field parameters were used in all simulations; key protein-ligand/$Mg^{2+}$ contacts observed in the crystal structure were closely monitored during simulations for validating the force field parameters. The chemical structure of the ligand is shown in Scheme S1, and the atom types and partial charges for the ligand are summarized in Table S3.

**Molecular dynamics data analysis.** CHARMM v41.0 was used to remove the overall translation and rotation and to orient all frames against the crystal structure before any analysis was performed. VMD and Pymol were used to visualize the trajectories. MDAnalysis, numpy, and scikit-learn packages were used for post-processing such as data analysis and plotting.

Correlated motions of protein residues are characterized with the covariance matrix of Cα carbons, which can be represented by either an N x N or 3N x 3N covariance matrix:

$$\mathbf{C} = <(\boldsymbol{q} - <\boldsymbol{q}>)(\boldsymbol{q} - <\boldsymbol{q}>)^T>$$

In the N x N matrix, $\boldsymbol{q} = (\boldsymbol{r}_1; \boldsymbol{r}_2; \ldots; \boldsymbol{r}_N)$ in which $\boldsymbol{r}_i = (x_i, y_i, z_i)$ is the three-dimensional atomic position of the ith atom. In the 3N x 3N matrix, $\boldsymbol{q} = (x_1, y_1, z_1, x_2, y_2, z_2, \ldots, x_N, y_N, z_N)^T$. After normalization, the covariance $C_{ij}$ ranges from -1 to 1. The N x N matrix is plotted in Fig. S7 for illustration, and to help highlight long-range correlated motions, the covariance of contact pairs is set to be 0; the contact probability of Cα pairs is defined as the fraction of frames in which the distance between the pair is less than 10 Å, and only pairs with a probability larger than 0.65 are recognized as contacts.

The 3N x 3N covariance matrix, which reveals more complex correlations, was used in the principal component analysis (PCA) without any filtering. The diagonalization of the 3N x 3N covariance matrix resulted in 3N eigenvalues, which are sorted in the descending order, as are the corresponding 3N eigenvectors.

To compare the difference in free energy landscapes and ensure the consistency in the direction of principal components among the different TetR variants, we merged the three sets of trajectories together and aligned all frames against the crystal structure of the wild type protein. PCA was carried out over the combined trajectory. The projection of each frame onto the eigenvectors resulted in 'Principal Components' (PCs), $V_i$. The first two PCs, $V_1$ and $V_2$, can be used to construct the two-dimensional free energy landscape:

$$\Delta G(V_1, V_2) = -k_B T[ln\rho(V_1, V_2) - ln\rho_{max}],$$

where $\rho(V_1, V_2)$ is an estimate of the joint probability density function obtained from the 2D histogram of the data. $\rho_{max}$ is the maximum density, which is subtracted to ensure $\Delta G = 0$ for the free energy minimum. The 1D free energy landscape can also be constructed as follows:

$$\Delta G(V_i) = -k_B T[ln\rho(V_i) - ln\rho_{max}].$$

The community analysis was carried out by the NetworkView plugin (*5*) in VMD with default setting. The definition of network is the same as that in the paper of Sethi et al (*6*). Each amino acid residue is represented by a node which is connected by edges. The edge weight is $w = -\log|C_{ij}|$, where $C_{ij}$ is the correlation between node i and node j. Here, the N x N covariance matrix was used in which both self-correlations and correlations with the nearest neighboring residues were set to be 0. The path distances between node i and node j are the sum of edge weights along the paths. The shortest distance between node i and node j is found by using the Floyd-Warshall algorithm. The Girvan-Newman algorithm (*7*) was used to partition the communities. Optimal communities can be found by maximizing the modularity value, Q, a measure of the difference in the probability of intra- and inter- community edge (*8*). Suboptimal path analysis was also done in NetworkView with a LengthOffset of 5. The sources are residues interacting with the ligand or $Mg^{2+}$ (residue id 64, 82, 100, 103, 116, 147) and the targets are residues directly involved in DNA-binding (residue id 26, 27, 28, 37, 38, 39, 40, 42, 43, 44, 48). For a given system, the occurrence of a residue is the number of times that the particular residue occurred in suboptimal paths below a threshold (i.e. the shortest distance + LengthOffset).

The quasi-harmonic approximation is commonly assumed in the calculation of configurational entropy of macromolecules. In our case, however, this approximation is unlikely to hold for at least the first two PCs, which exhibit very anharmonic landscape as illustrated in Fig. 4 in the main text. Therefore, the PC1&2 and other PCs were treated differently in the calculation of configurational entropies. In short, we followed the same procedure as in the paper of Andricioaei et al (*43*). except that the PC1&2 were excluded. Then, the contribution of PC1&2 were calculated separately as follows:

$$S = k\ln\frac{\sqrt{2\pi k_B T}}{h}\sum_i e^{-\beta U_i} + \frac{k}{2} + \frac{\sum_i U_i e^{-\beta U_i}}{T\sum_i e^{-\beta U_i}},$$

in which $k_B$ is the Boltzmann constant, $T$ is the absolute temperature, $h$ is the Planck constant, $\beta = 1/k_B T$, and $U$ is the effective potential along PC1 or PC2. The summation index $i$ runs over the number of bins in the corresponding 1D histogram. Since the number of alpha carbons in the wild type and mutants are the same, we can directly compare the difference in configurational entropy between the wild type and each mutant:

$$T\Delta S = TS_{mutnat} - TS_{wild\ type}$$

We plot the time evolution of $T\Delta S$ along the simulated trajectories in Fig. S8A; each point in the curves represents results that include all frames up to that time.

To demonstrate the robustness of the landscape comparison between different TetR variants, the free energy landscapes were also analyzed using the locally scaled diffusion map (LSDmap) (*21*, *22*). LSDmap uses a Gaussian kernel to describe the transition probability between two conformations,

$$K_{ij} = \exp\left(-\frac{||\boldsymbol{x}_i - \boldsymbol{x}_j||^2}{2\varepsilon_i\varepsilon_j}\right),$$

where $K_{ij}$ is the transition probability, $||\mathbf{x}_i - \mathbf{x}_j||^2$ is the RMSD between two conformations, $\varepsilon_i$ and $\varepsilon_j$ are the local scales of the corresponding conformations. We used the procedure proposed in literature (*21*) to determine the local scales. $K_{ij}$ represents the ability of the diffusion of one conformation to the other. This matrix can be easily converted to a Markov matrix, whose eigenvectors represent the diffuse coordinates (DCs). We projected the conformations onto the first two DCs to obtain the corresponding free energy landscapes.

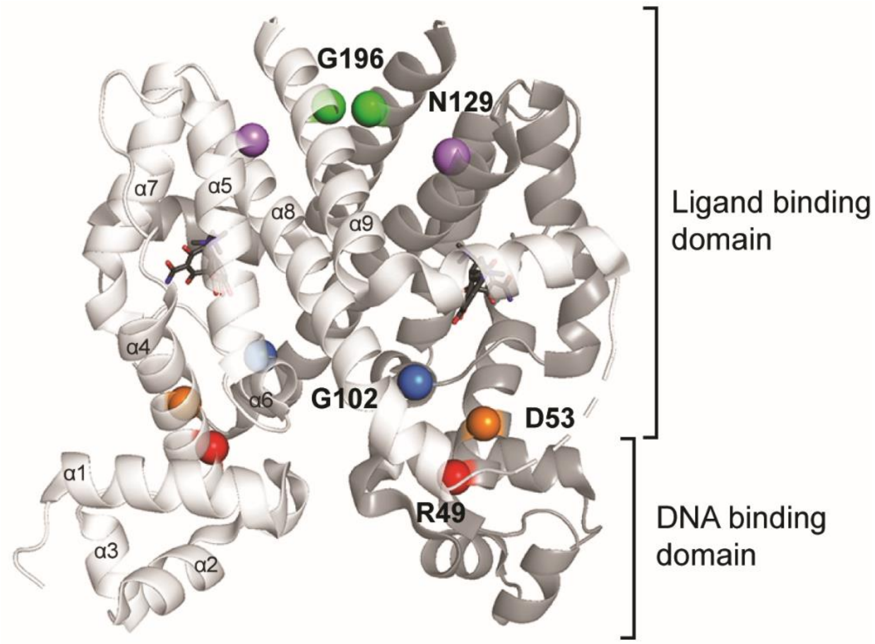


**Fig. S1. TetRb structure.**

Crystal structure of TetR(B) with bound [Minocycline:Mg]$^{+}$ (PDB ID: 4ac0). Dead variants are labeled as colored spheres on both monomers of the TetR dimer. The ligand and DNA binding domains of the dimer are indicated with alpha helices numbered on one monomer.

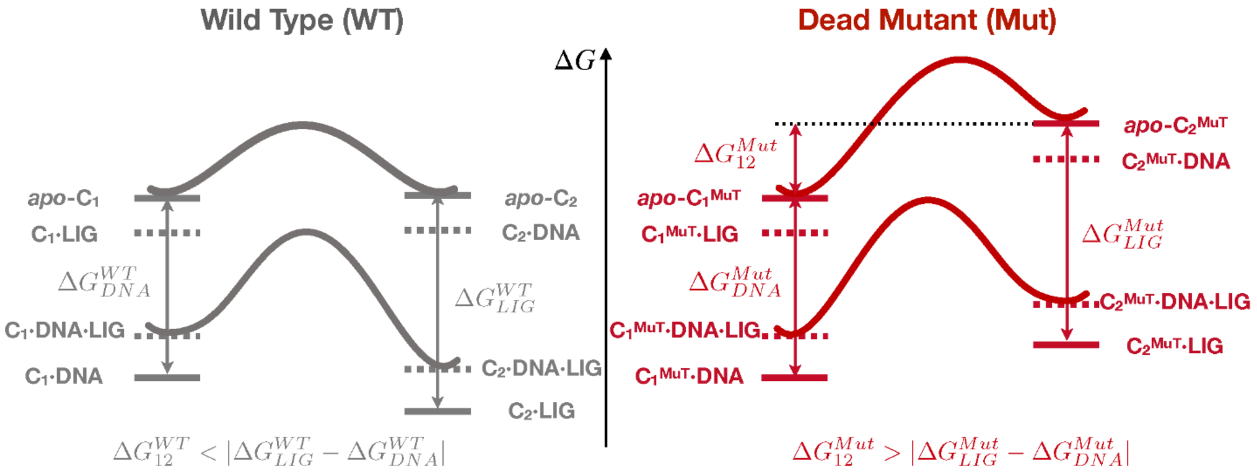


**Fig. S2. Comparison of detailed schematic free energy levels and landscapes for the wild type and a dead mutant (Mut) of TetR.**

In this simple model, each protein is assumed to have two conformational states ($C_1/C_2$), which preferably bind to the DNA and the inducer ligand, respectively; for simplicity, we do not separately consider the two ligand/DNA binding domains in each dimeric TetR. Since each protein can adopt four chemical states: apo, the protein-DNA binary complex, the protein-ligand binary complex and the protein-DNA-ligand ternary complex, there are eight stereochemical states for each protein. The effective free energy levels (which in general depend also on the bulk concentrations of ligand and DNA) of these states are indicated with horizontal bars, with the stereochemical states expected to have very low populations shown in dashed bars. The free energy landscape is well defined for systems of the same molecular composition and only two examples for each protein are shown for clarity. In the wildtype protein, the binding affinity of the ligand to apo-$C_2$ is larger in magnitude than that of the DNA to apo-$C_1$; assuming that apo-$C_1$ and apo-$C_2$ are similar in free energy, this model predicts $C_2$·ligand as the predominant species (i.e., inducer binding leads to dissociation from the DNA). In a dead mutant, with a simple model, the intrinsic binding affinities of $C_1^{Mut}$ to DNA and $C_2^{Mut}$ to ligand are not perturbed relative to wildtype, but apo-$C_2^{Mut}$ is destabilized relative to apo-$C_1^{Mut}$ by an amount of $\Delta G_{12}^{Mut}$; if $\Delta G_{12}^{Mut}$ is larger in magnitude than the differential DNA/ligand binding affinity, $\Delta G_{Lig}^{Mut} - \Delta G_{DNA}^{Mut}$, the model predicts that $C_1^{Mut}$·DNA is the predominant population even in the presence of the inducer ligand. In other words, $\Delta G_{12}^{Mut}$ is the energetic difference that abolishes ligand inducibility by destabilizing the active ($C_2^{Mut}$) state (or, equivalently, stabilizing the inactive state, $C_1^{Mut}$). On the other hand, if $\Delta G_{12}^{Mut}$ is smaller in magnitude than $\Delta G_{Lig}^{Mut} - \Delta G_{DNA}^{Mut}$, $C_2^{Mut}$·ligand is still the predominant species; i.e., reduction of $\Delta G_{12}^{Mut}$ is likely the mechanism for rescued mutant.

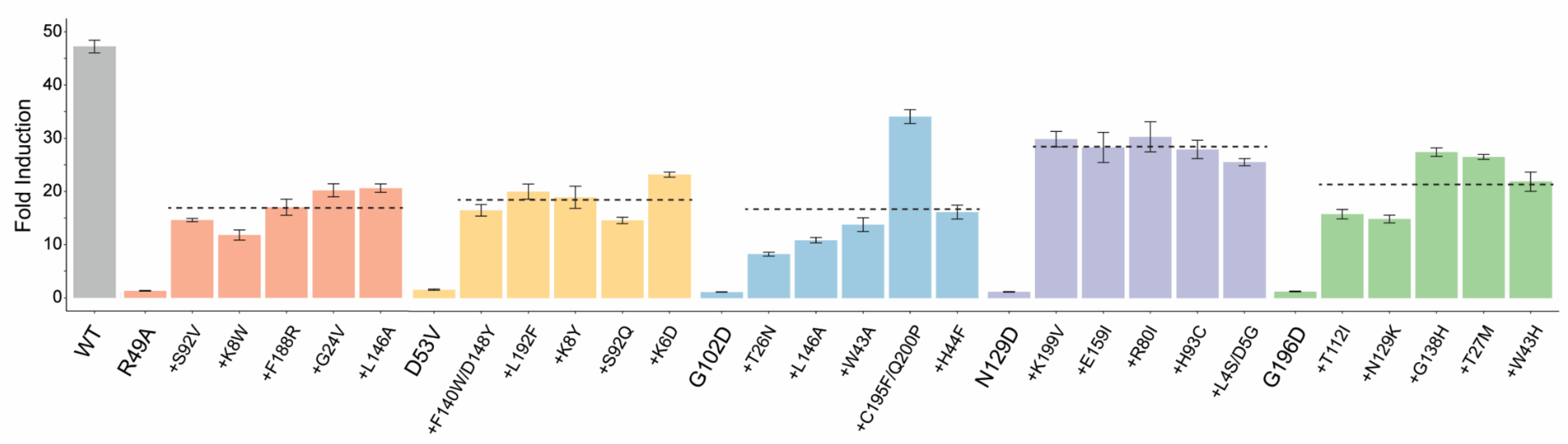


**Fig. S3. Fold induction of dead and rescue variants.**

Fold induction of wildtype, dead, and rescued variants is shown as ratio of fluorescence in the induced and uninduced states. Average fold induction of rescued variants for each dead variant is shown as dashed lines. Error bars represent standard error of three biological replicates.

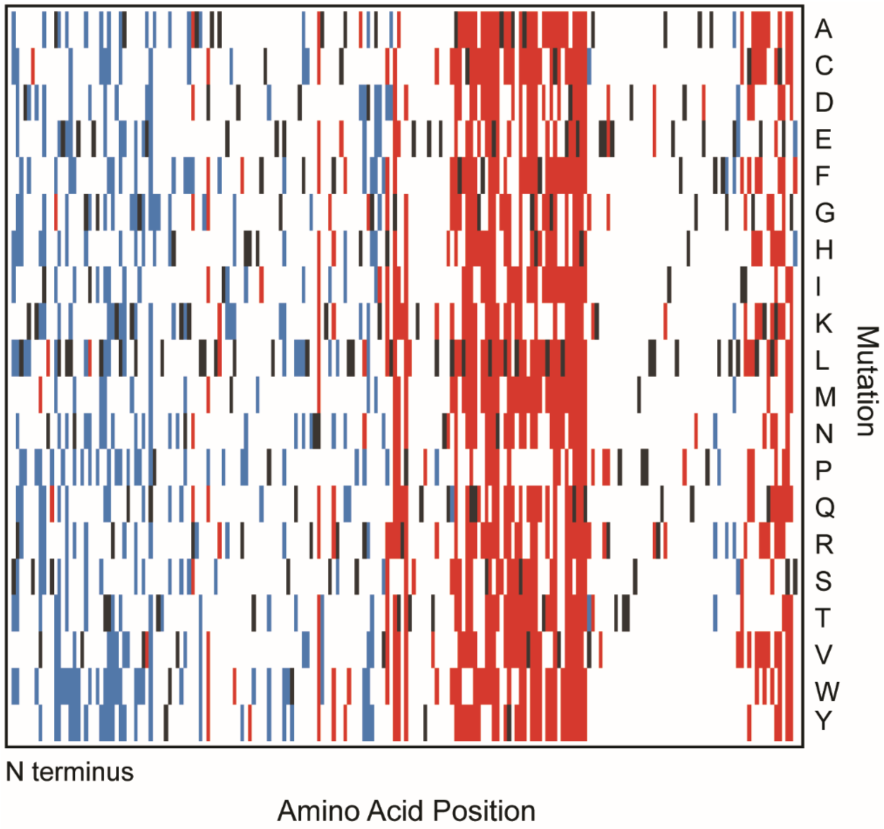


**Fig. S4. Dead and broken mutations in TetR.**

Dead and broken variants are shown in red and blue, respectively, with the wildtype residue indicated in black.

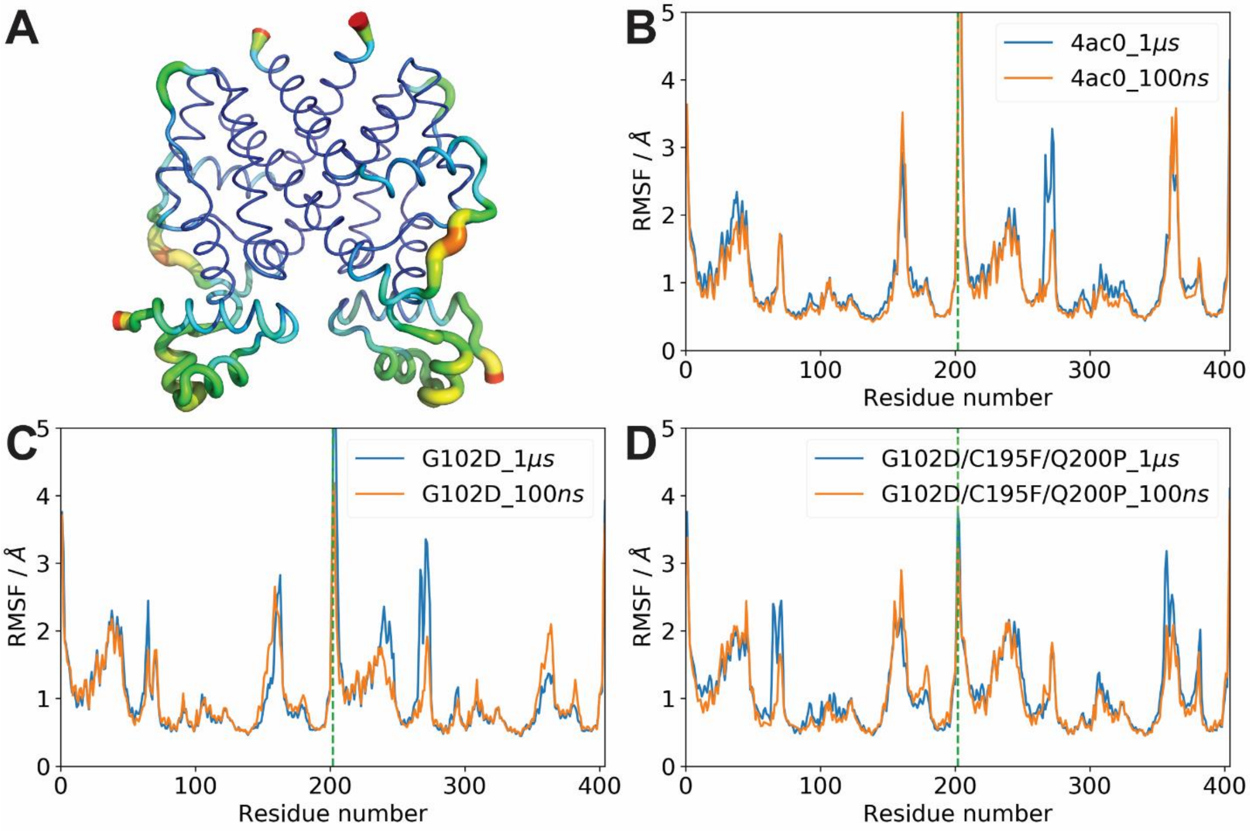


**Fig. S5. Structural stability and residue flexibility of TetR in MD simulations.**

(**A**) Average structure of the wild type protein with the thickness and color indicating the magnitude of RMSF (root mean square fluctuation). (**B-D**) RMSF of the three systems averaged over 100 ns (orange) and 1 μs (blue).

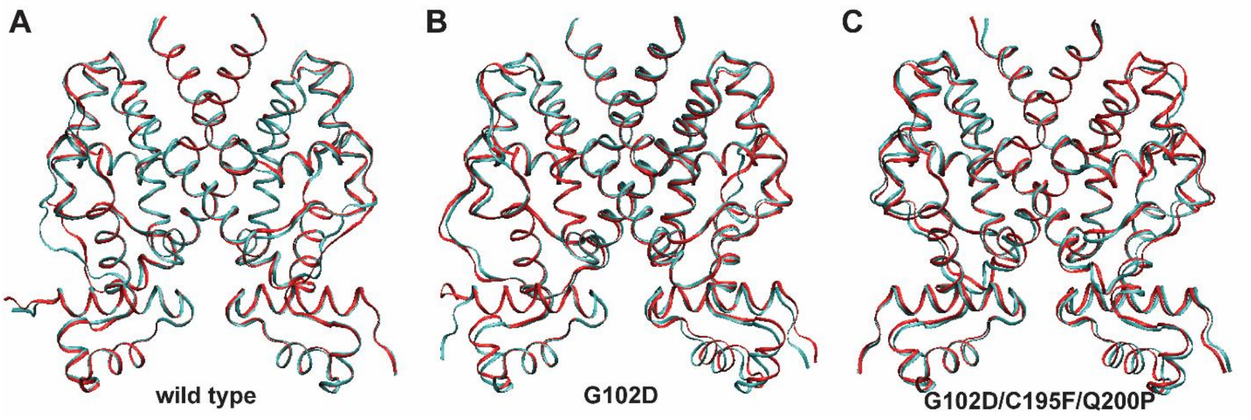


**Fig. S6. Convergence and comparison of average structures.**
(**A-C**) Overlaid average structure of 100 ns (red) and 1 μs (cyan).

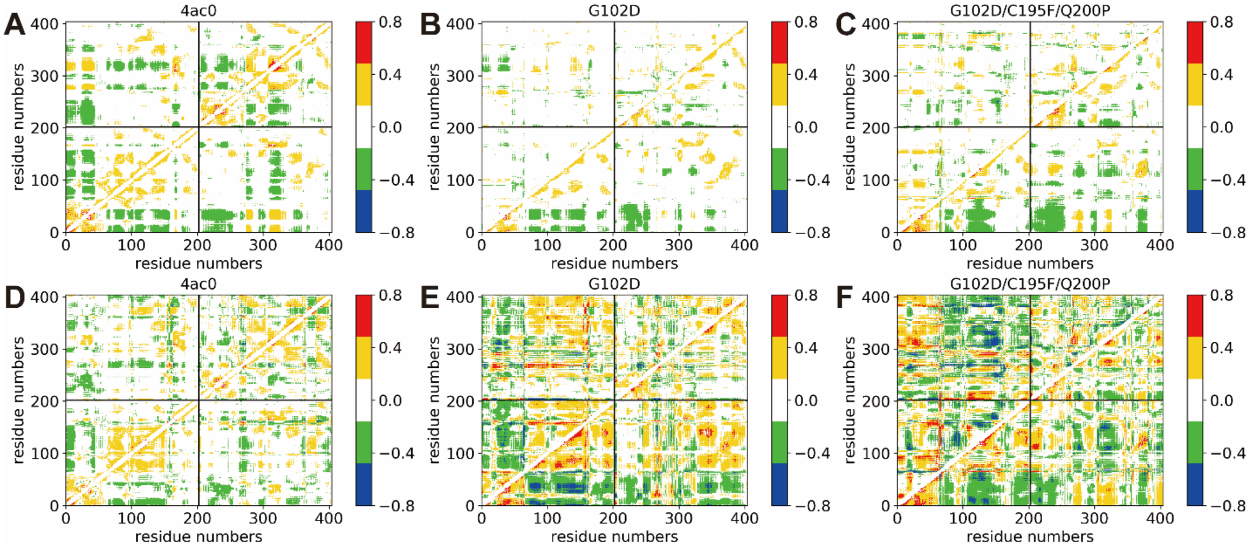


**Fig. S7. Covariance of Cα atoms and the convergence of the covariance matrix for wildtype, G102D, and G102D/C195F/Q200P.**

(**A-C**) The covariance matrix averaged over the 1 μs trajectories; the upper triangles in (**B**) and (**C**) show the difference in covariance between mutant and wild type proteins. (**D-F**) The covariance matrix averaged over 100 ns trajectories; the upper triangles in (**E**) and (**F**) show the difference in covariance between mutant and wild type proteins.

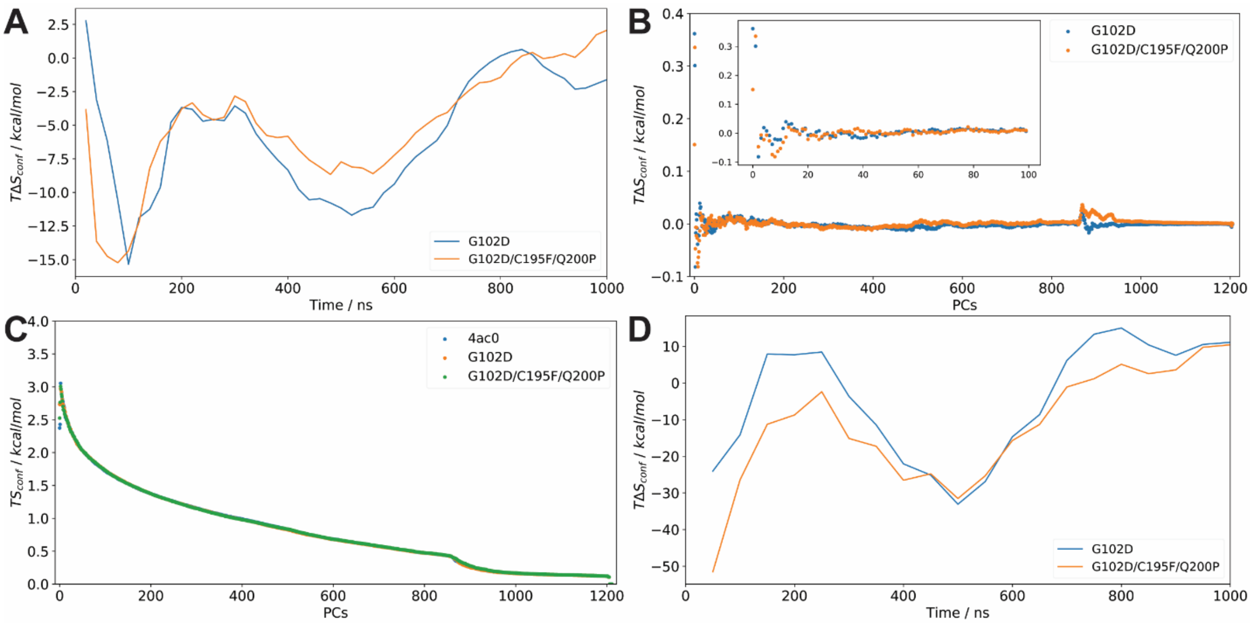


**Fig. S8. Conformational entropy converges slowly.**

Alpha carbon atoms (**A-C**) in protein were used to calculate the configurational entropy using the modified quasi-harmonic analysis (see text). The temperature, $T$, is 303.15 K. (**A**) $\Delta S = S_{mutant} - S_{wild\ type}$ as a function of simulation time for the two mutants relative to the wild type protein; (**B**) entropic contribution of each mode relative to wild type using the complete (1□s) trajectories; (**C**) entropic contribution of each mode using the complete (1□s) trajectories; (**D**) all heavy (non-hydrogen) atoms were also used to calculate configurational entropy using the same method as above.

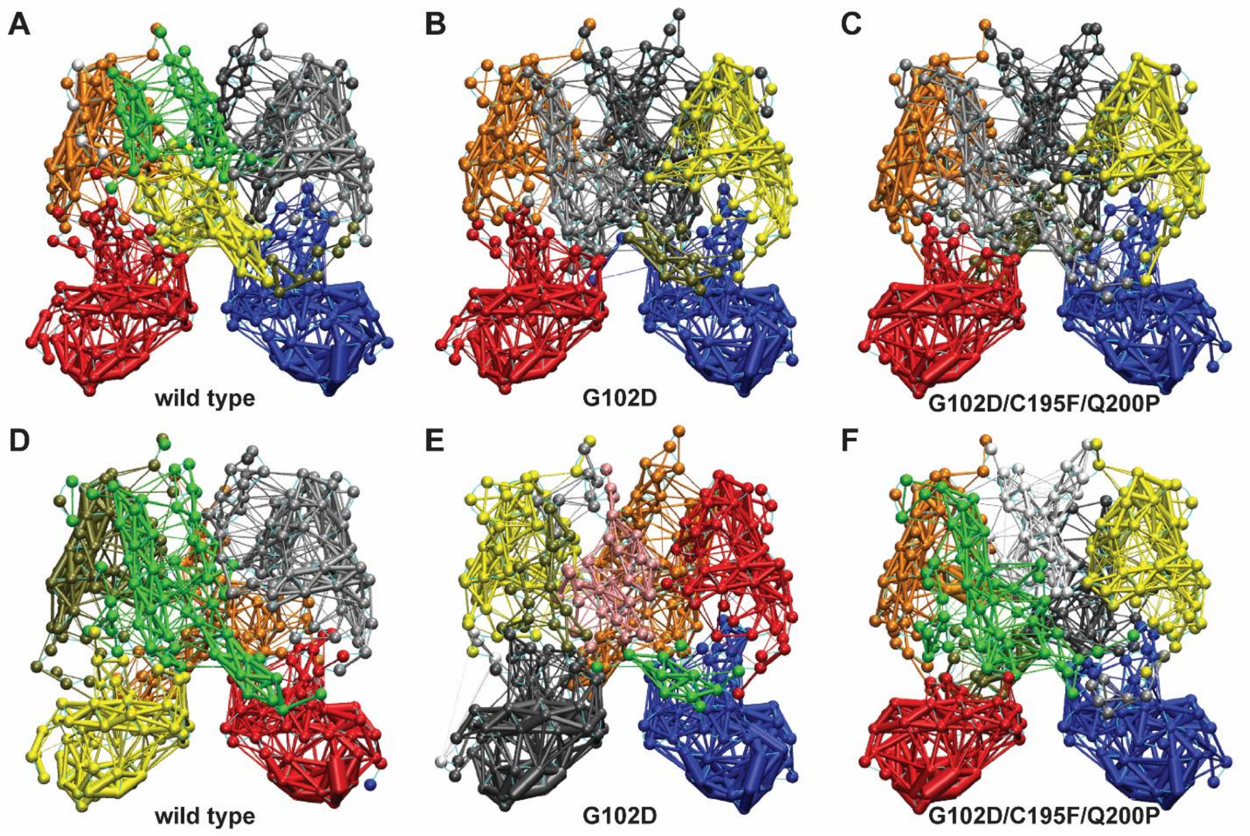


**Fig. S9. The partition of structure into communities and convergence of the analysis.**
(**A-C**) The communities of each system obtained from the 1 μs trajectories. (**D-F**) The communities of each system obtained from the first 100 ns trajectories. Each color represents one community and the choice of color is arbitrary. The solid spheres (nodes) represent alpha carbons and the thickness of lines connecting two nodes represent the magnitude of correlation between two nodes.

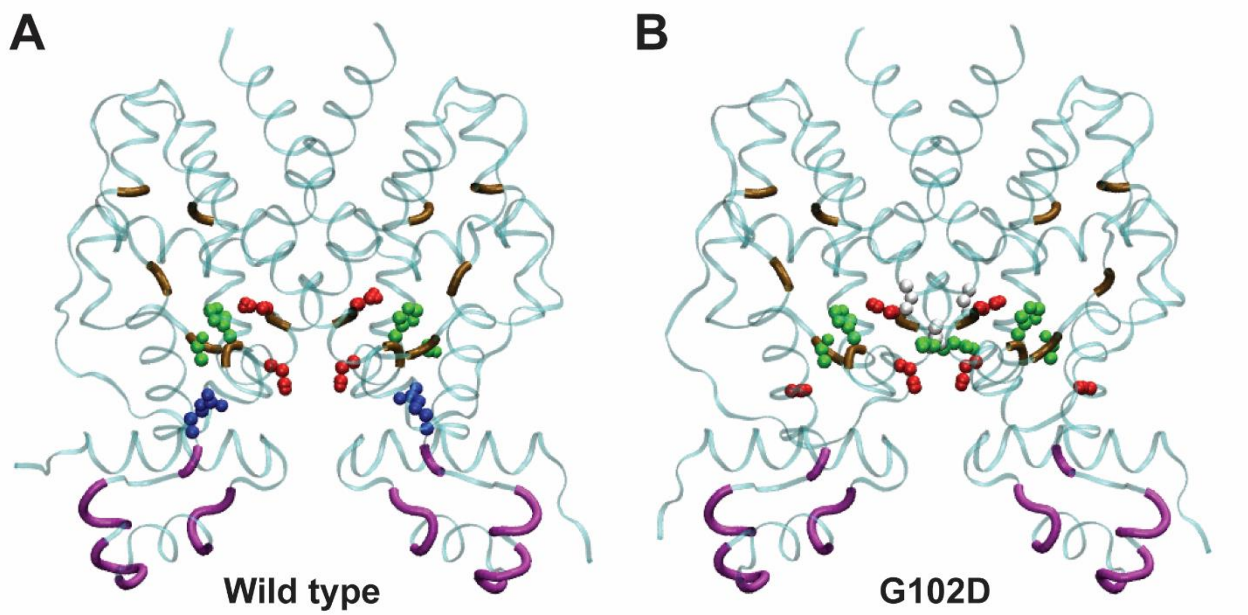


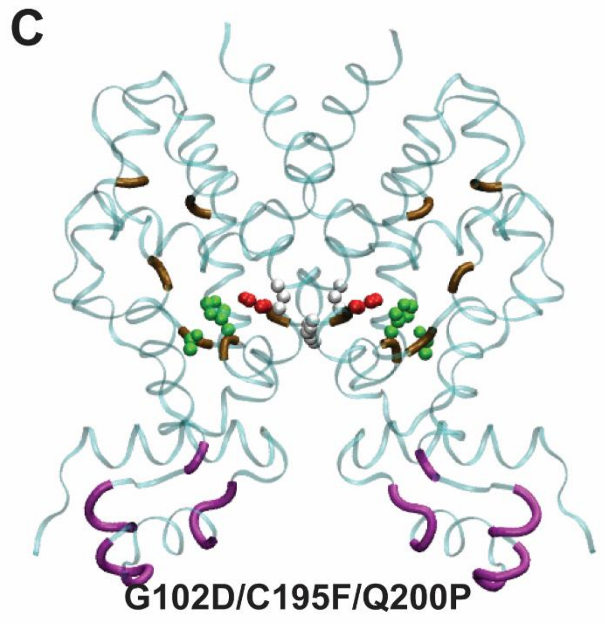


D

| | Suboptimal path | Deads in experiment |
|---|---|---|
| W.T. | 10.0% | 9.1% |
| G102D | 16.0% | 14.5% |
| G102D/ C195F/ Q200P | 10.0% | 9.1% |

**Fig. S10. Overlap between 'hotspot' residues identified in experiments and 'hub' residues found in suboptimal path analysis using MD trajectories.**

(**A-C**) The protein structures are represented by ribbons and colored by residue types (white for non-polar residues, green for polar residues, blue for negatively-charged residues, and red for positively-charged residues). Residues in ligand-binding domain and residues directly bound to DNA are represented by tubes. The overlapped residues were highlighted as van der Waals spheres in the structures. (**D**) Hub residues are compared with the dead variants in experiments. Hub residues are defined as the top 50 most occurred residues in suboptimal paths for a given TetR variant. Percentages in the second and third columns are the fractions of common residues among the hub residues and the deads in experiments, respectively.

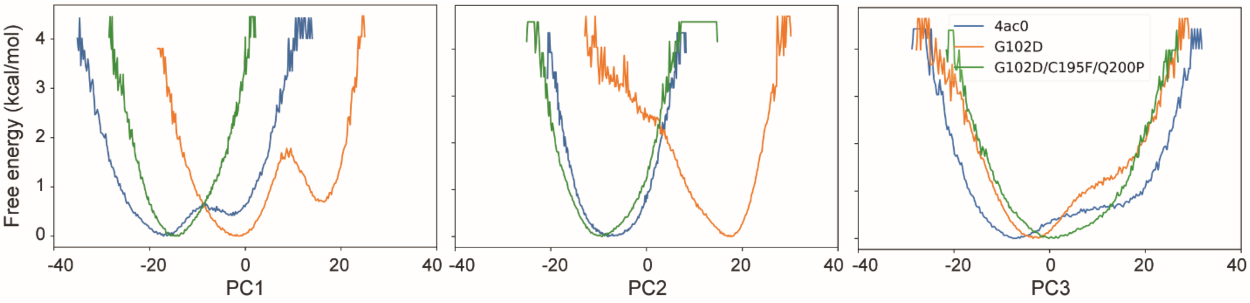


**Fig. S11. One-dimensional free energy landscapes along the first three principal components.**

Along PC1 and PC2, the landscapes show convergence of the wildtype and rescued variant, but not the dead variant. Beyond PC1 and PC2, the landscapes of all three systems show little difference, indicating that the first two principal components capture the most difference in the free energy landscape.

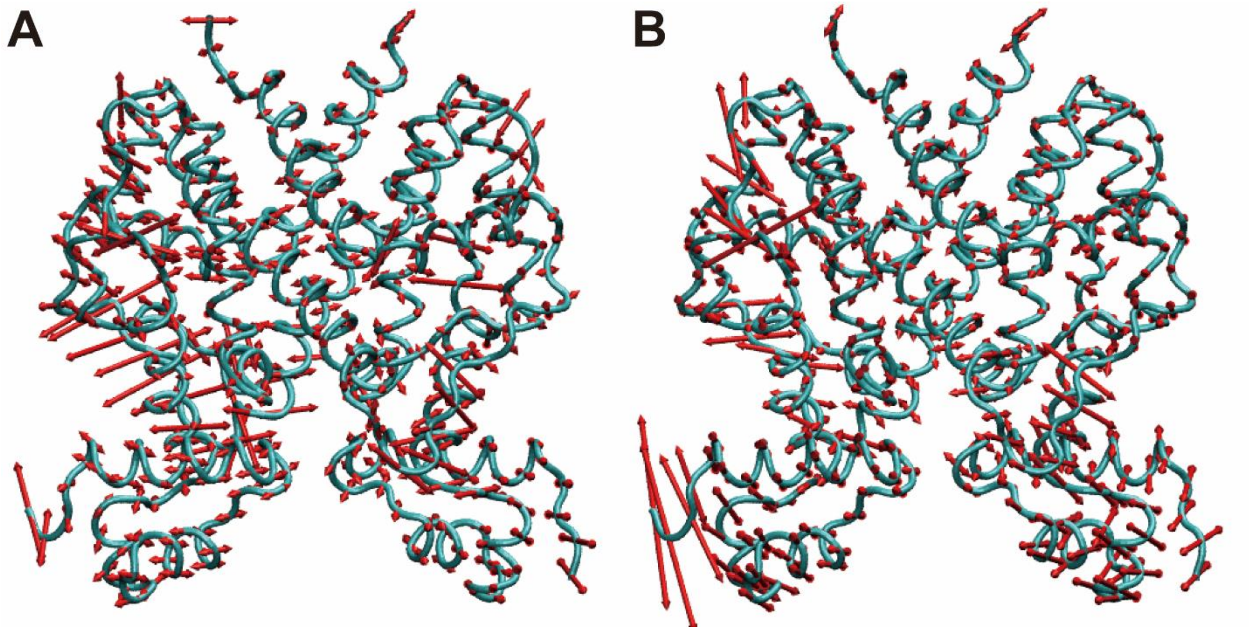


C

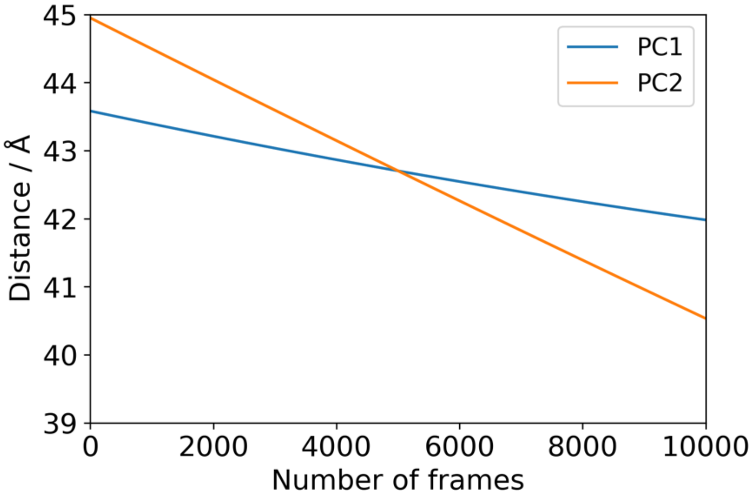


**Fig. S12. Directionality and relative magnitude of motions along the first two principal components (also see Movies S1 and S2).**

(**A**) First principal component (PC1) mainly represents the motion of loops. (**B**) Second principal component (PC2) represents motions of both loops and DNA-binding domains. The red arrows show the direction of alpha carbons in protein and the length of arrows show the relative magnitude of motion. The pendulum type of motions of the DNA binding domains were proposed to affect the DNA binding affinity and thus activity of TetR. (**C**) As an example for the evolution of key structural features involved in the first two principal components, the distance between the two □3 helices as a function of displacement along PC1/PC2 is plotted. The plot highlights that both principal components involve relative displacement of the DNA binding domains, a structural transition that has been proposed to modulate the DNA binding activity (*23*, *24*)

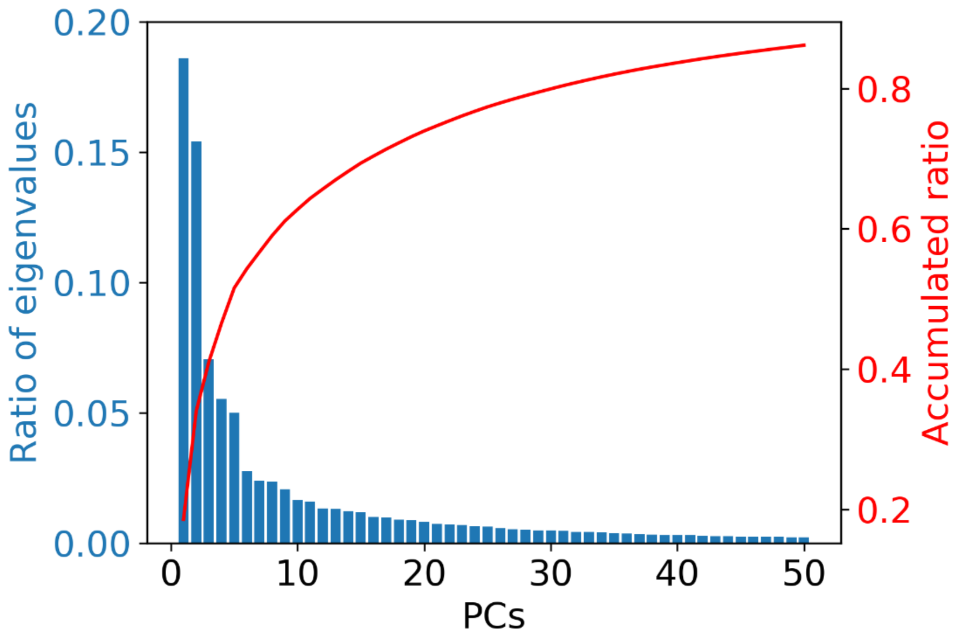


**Fig. S13. The first two principal components represent a significant fraction of the motions.** The eigenvalues were obtained after performing PCA on the combined trajectories of wild type, G102D, and G102D/C195F/Q200P. Normalization of eigenvalues resulted in the ratios. Although TetR is structurally rather rigid, the first sets of principal components capture a significant fraction of the overall motion; this justifies the use of the first few leading principal components in the free energy landscape analysis discussed in the main text.

| Dead Variant | Compensatory Mutations |
|---|---|
| **R49A** | D5AL, K6HW, G24A, K29IS, E37Y, R87K, H93W, T112P, P184K |
| **D53V** | R3V, L4Y, D5A, K6N, K8H, I22C, G24A, L25I, G35DN, R94W, L113C, P184T |
| **G102D** | L146A |
| **N129D** | L4TY, D5V, K6R, S7P, K8DNW, V9G, A13C, L14N, E19D, I22V, K29Q, L34F, G35F, V36I, E37FH, Q38F, W43K, V45C, K46Y, N47H, L52T, A54FN, A56Q, I57L, E58Y, L60E, H63S, H66Y, F67Y, C68M, P69H, E71L, N81AG, A83GN, T84IT, S85CW, F86CEM, R87Q, C88EHM, A89KY, L91HM, S92M, A97M, K98E, V99F, H100AT, L101H, G102H, R104CSVY, T106V, E107RW, K108S, Q109I, Y110L, E111AIY, T112HNP, L113S, T163L, D164Y, A173E, Q180I, P184V, F188G, I193AC, I184K, C195KTV, G196DIMNPQ, L197AFHPV, E198R, K199CFLQTWY, Q200GHKV, L201GIK, K202ENV, C203DIQSTY, E204ILQST, S205GMVWY, G206CHLMT, S207PQ |
| **G196D** | S2CD, D5T, T27M, V36F, E37G, P39S, W43HY, V45FY, A50E, D53V, I57HQ, H63Y, F67CDKLQRSV, C68L, C88R, E107Q, T112IMQRV, F119G, C121P, L131AIVY, S135M, G138HNQ, H139S, L146T, V153R |

**Table S1. All single-mutant compensatory mutations identified for each dead variant.**

| TetR Library | | Total reads | Good reads | Translated reads | Single mutant reads | Number of variants | Number of variants over 10x threshold | Coverage |
|---|---|---|---|---|---|---|---|---|
| Replicate 1 | Presorted | 542436 | 440570 | 389212 | 272093 | 3759 | 3442 | 88% |
| | aTC- | 624025 | 509408 | 477335 | 349355 | 3603 | 2981 | |
| | aTC+ | 651400 | 579330 | 484690 | 320675 | 1912 | 1213 | |
| Replicate 2 | Presorted | 654463 | 525320 | 468679 | 336478 | 3774 | 3456 | 88% |
| | aTC- | 642040 | 562346 | 520758 | 380410 | 3611 | 3005 | |
| | aTC+ | 682668 | 597205 | 496071 | 336546 | 2027 | 1249 | |

**Table S2. Next-generation sequencing statistics of the single-mutant TetR library.**

| Atom name | Atom type | Partial charge | Atom name | Atom type | Partial charge |
|---|---|---|---|---|---|
| **O1** | OG2D3 | -0.48 | **O6** | OG311 | -0.39 |
| **C1** | CG2O5 | 0.442 | **C21** | CG2DC1 | 0.18 |
| **C2** | CG2R61 | -0.331 | **C22** | CG2O1 | 0.567 |
| **C3** | CG2R61 | 0.407 | **O7** | OG2D1 | -0.514 |
| **O2** | OG312 | -0.762 | **N2** | NG2S2 | -0.714 |
| **C4** | CG2R61 | -0.32 | **H1** | HGP1 | 0.42 |
| **C5** | CG2R61 | 0.107 | **H2** | HGR61 | 0.115 |
| **O3** | OG311 | -0.53 | **H3** | HGR61 | 0.115 |
| **C6** | CG2R61 | -0.116 | **H4** | HGR61 | 0.115 |
| **C7** | CG2R61 | -0.113 | **H5** | HGA3 | 0.09 |
| **C8** | CG2R61 | -0.117 | **H6** | HGA3 | 0.09 |
| **C9** | CG2R61 | 0.008 | **H7** | HGA3 | 0.09 |
| **C10** | CG2R61 | -0.002 | **H8** | HGA2 | 0.09 |
| **C11** | CG331 | -0.266 | **H9** | HGA2 | 0.09 |
| **C12** | CG2R61 | -0.012 | **H10** | HGA1 | 0.09 |
| **C13** | CG321 | -0.183 | **H11** | HGP1 | 0.42 |
| **C14** | CG301 | 0.405 | **H12** | HGA1 | 0.09 |
| **C15** | CG2O5 | 0.323 | **H13** | HGA3 | 0.09 |
| **O4** | OG2D3 | -0.471 | **H14** | HGA3 | 0.09 |
| **O5** | OG311 | -0.714 | **H15** | HGA3 | 0.09 |
| **C16** | CG311 | -0.071 | **H16** | HGA3 | 0.09 |
| **C17** | CG311 | 0.091 | **H17** | HGA3 | 0.09 |
| **N1** | NG301 | -0.616 | **H18** | HGA3 | 0.09 |
| **C18** | CG331 | -0.099 | **H19** | HGP1 | 0.42 |
| **C19** | CG331 | -0.099 | **H20** | HGP1 | 0.29 |
| **C20** | CG2D1O | 0.035 | **H21** | HGP1 | 0.29 |

**Table S3. The atom name, atom type, and partial charge for the ligand**

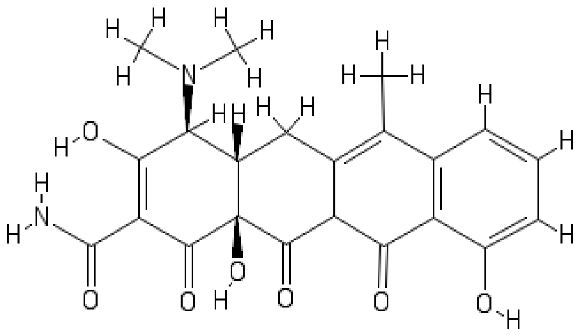


**Scheme S1. Chemical structure of anhydrotetracycline**

**Movie S1. The motion of protein along the first principal component**

The motion with the largest amplitude occurred in the loop region, which is consistent with previous analysis. The DNA-binding domain (DBD) resembled the 'pendulum-like' motion, while the patterns of motion are different in two PCs (also see Movie. S2). In PC1, the DBD in two monomers moved toward the same direction.

**Movie S2. The motion of protein along the second principal component**

The motion with the largest amplitude occurred in the loop region, which is consistent with previous analysis. The DNA-binding domain (DBD) resembled the 'pendulum-like' motion, while the patterns of motion are different in two PCs (also see Movie. S1). In PC2, the DBD in two monomers moved toward the opposite direction.

**Dataset S1. Phenotypic summary of all TetR mutations (separate file)**

A matrix of all mutations found to be dead (D) or broken (B) indicated at each position in the protein along with the wildtype residue (WT). Blank cells did not appear in the library or could not be definitively classified as either dead or broken. Positions with five or more mutations that inactivate or break the protein were labeled with a Dead or Broken phenotype. The calculated conservation and centrality scores for each position are present.

**SI References**


1. I. Rivalta, M. M. Sultan, N.-S. Lee, G. A. Manley, J. P. Loria, V. S. Batista, Allosteric pathways in imidazole glycerol phosphate synthase. *Proc. Natl. Acad. Sci. USA* **109**, E1428-E1436 (2012).
2. J. Guo, X. Pang, H.-X. Zhou, Two pathways mediate interdomain allosteric regulation in Pin1. *Structure* **23**, 237-247 (2015).
3. B. Chakrabarty, N. Parekh, NAPS: Network analysis of protein structures. *Nucleic Acids Res.* **44**, W375-W382 (2016).
4. A. Sethi, J. Eargle, A. A. Black, Z. Luthey-Schulten, Dynamical networks in tRNA:protein complexes. *Proc. Natl. Acad. Sci. USA* **106**, 6620-6625 (2009).
5. M. Girvan, M. E. J. Newman, Community structure in social and biological networks. *Proc. Natl. Acad. Sci. USA* **99**, 7821-7826 (2002).
6. M. E. J. Newman, Modularity and community structure in networks. *Proc. Natl. Acad. Sci. USA* **103**, 8577-8582 (2006).
7. I. Andricioaei, M. Karplus, On the calculation of entropy from covariance matrices of the atomic fluctuations. *J. chem. Phys.* **115**, 6289-6292 (2001).
